# Eleven deep-sea coral genome assemblies unveil insights into evolution, adaptation, and coral biodiversity

**DOI:** 10.64898/2026.05.06.723128

**Authors:** Nannan Zhang, Liangwei Li, Kaiwen Ta, Chengcheng Shi, Inge Seim, Yingying Zhang, Weijia Zhang, Zhen Cui, Xueyan Xiang, Limin Jia, Qijin Ge, Mengran Du, Tongtong Xie, Qianyue Ji, Zhen Yue, Guangyi Fan, Shanshan Liu, Liang Meng

**Affiliations:** BGI Research, Sanya 572025, China; BGI Research, Qingdao 266555, China; Institute of Deep-seas Science and Engineering, Chinese Academy of Sciences, Sanya 572000, China; BGI Research, Shenzhen 518083, China; College of Life Sciences, University of Chinese Academy of Sciences, Beijing 100049, China; MGI Tech, Shenzhen 518083, China; Hainan Technology Innovation Center for Marine Biological Resources Utilization(Preparatory Period), BGI Research, Sanya 572025, China; State Key Laboratory of Genome and Multi-omics Technologies, BGI Research, Shenzhen 518083, China; Shenzhen Key Laboratory of Marine Biology Genomics, BGI Research, Shenzhen 518083, China; Institution of Deep-Sea Life Sciences, IDSSE-BGI, Hainan Deep-sea Technology Laboratory, Sanya 572000, China

## Abstract

Deep-sea corals are vital in maintaining coral ecosystem biodiversity, yet their genetic characteristics remain largely unexplored. Here, we present 11 deep-sea coral genome assemblies, including four Hexacorallia and seven Octocorallia species, significantly contributing new genomic information across two orders. Our analysis reveals the historical dynamics of coral speciation and the influence of environmental factors on the evolution of coral reef ecosystems.Total of 126 horizontal gene transfer (HGT) events were detected, among which genes from the ancestor of symbiodiniaceae indicate that the ancestors of deep-sea corals may have inhabited shallow-sea environments. Notably, several of these HGTs are involved in phosphorus (*PhnX*/*PhnW*) and cholesterol (*DHCR7*) metabolisms within corals, indicating that HGTs may serve as an adaptive survival strategy for the coral holobionts. Deep-sea corals also rely on symbiotic bacteria to synthesize 10 essential amino acids (such as valine and tyrosine), retaining only partial amino acid synthesis capacity. In addition, we investigated the evolution of key biological rhythm genes and temperature adaptation in corals. The loss of key rhythm genes (e.g., *clock* and *cry*) in deep-sea corals and copy number difference of genes related to heat stress (e.g., *Cbl-b* and *Rchy*) revealed genetic difference between deep-sea and shallow-sea corals. Our new genome assemblies enhance the understanding of deep-sea coral evolution, biodiversity, and adaptation, providing a genetic foundation for coral conservation.

## Introduction

Corals have a global distribution and are commonly found along the edges of continental shelves and seamounts in oceans worldwide^1-3^. In addition to representing marine biodiversity, they also play a crucial role in the reproduction and survival of many marine species by supporting various fishes, invertebrates, and other types of corals^3,4^. These organisms rely on the structural complexity of corals to establish themselves and form communities^5-7^. Global temperature changes are one of the primary drivers of coral speciation^8-11^. The Earth’s climate has experienced significant fluctuations throughout its history, with periods of cooling and warming. These temperature changes profoundly affect coral populations, influencing their genetic diversity and ecological interactions. Rapid speciation in response to global temperature changes allows corals to occupy new ecological niches and expand their distribution, enhancing the overall diversity of coral reefs^12,13^.

Deep-sea corals thrive in the depths of the ocean floor, growing on rocky or sedimentary substrates at depths greater than 200 meters^14^. Despite being important and unique components of deep-sea ecosystems, deep-sea corals face challenges in adapting to harsh conditions, such as the absence of light and photosynthesis. They cannot rely on symbiotic Symbiodiniaceae for nutrition, unlike their shallow-sea counterparts. Instead, they primarily obtain food from organic matter that settles from the ocean surface^7,14^. Recent studies suggest that endosymbionts are a nutrient source for deep-sea corals^15^.

Corals form symbiotic relationships with various microorganisms^16,17^. They rely on symbiotic microorganisms to acquire metabolites crucial for their physiological processes, including amino acids, vitamins, and lipids^18-20^. Previous studies have shown that corals cannot synthesize several essential amino acids but acquire them from their symbionts^21-23^. For instance, in *Porites australiensis*, 11 amino acids might be provided by coral-associated microbes^23^. Some coral species cannot synthesize cholesterol and rely on their algae to acquire this essential compound^24^. Another mechanism for acquiring metabolites is horizontal gene transfer (HGT). HGT contributes to the acquisition of genes and metabolic pathways by corals ^16,25,26^. Previous studies highlight the transfer of genetic material from other organisms, such as bacteria and algae^25,26^, into coral genomes.

Genetic studies have provided valuable insights into the evolution of corals. Recent studies have reconstructed phylogenetic trees of a wide range of coral lineages using UCEs and exons ^27-29^, but more robust and reliable tree based on whole-genome data are limited. In addition, the origin of corals is a topic of scientific debate, with deep and shallow sea origin hypotheses proposed. Some studies on corals explored the origins of a particular lineage of corals through molecular datasets and fossil records and have come up with completely opposite results^30,31^. This is due to the different identification of the timepoint represented by “origin” and the different breadth of species sampling in degenerate analysis. To date, most published coral genomes are from shallow-sea corals^26,32-40^, which prevents more comprehensive evolutionary analysis.

Our understanding of deep-sea coral behavior and ecology has significantly improved by advancements in deep-sea exploration technology. However, a lack of knowledge remains on their genetics and evolution. In this study, we *de novo* assembled the genomes of 11 deep-sea coral species from two subclasses. By analyzing these genomes, we investigated the association between the rapid speciation of corals and global temperature changes. Additionally, we systemically compared the molecular mechanisms of the environmental adaptation between shallow-sea and deep-sea corals. Our findings provide insights into coral evolution, adaptations, and coral biodiversity.

## Results

### Eleven deep-sea coral genome features and phylogeny

We employed PacBio HiFi-CCS sequencing technology to generate 315.34 Gb of whole-genome sequencing data for 11 deep-sea corals (Figure 1A, Table S1 and 2). The genome assembly sizes ranged from 445.95 Mb to 1.24 Gb, with contig N50 values (428.50 Kb to 10.75 Mb) longer than most published coral genomes (Figure 1B). The average BUSCO (Benchmarking Universal Single-Copy Orthologs) completeness score of our 11 assemblies was also better than 92 other coral genomes (average BUSCO 79.3%) (Table S3).

**Figure 1.**
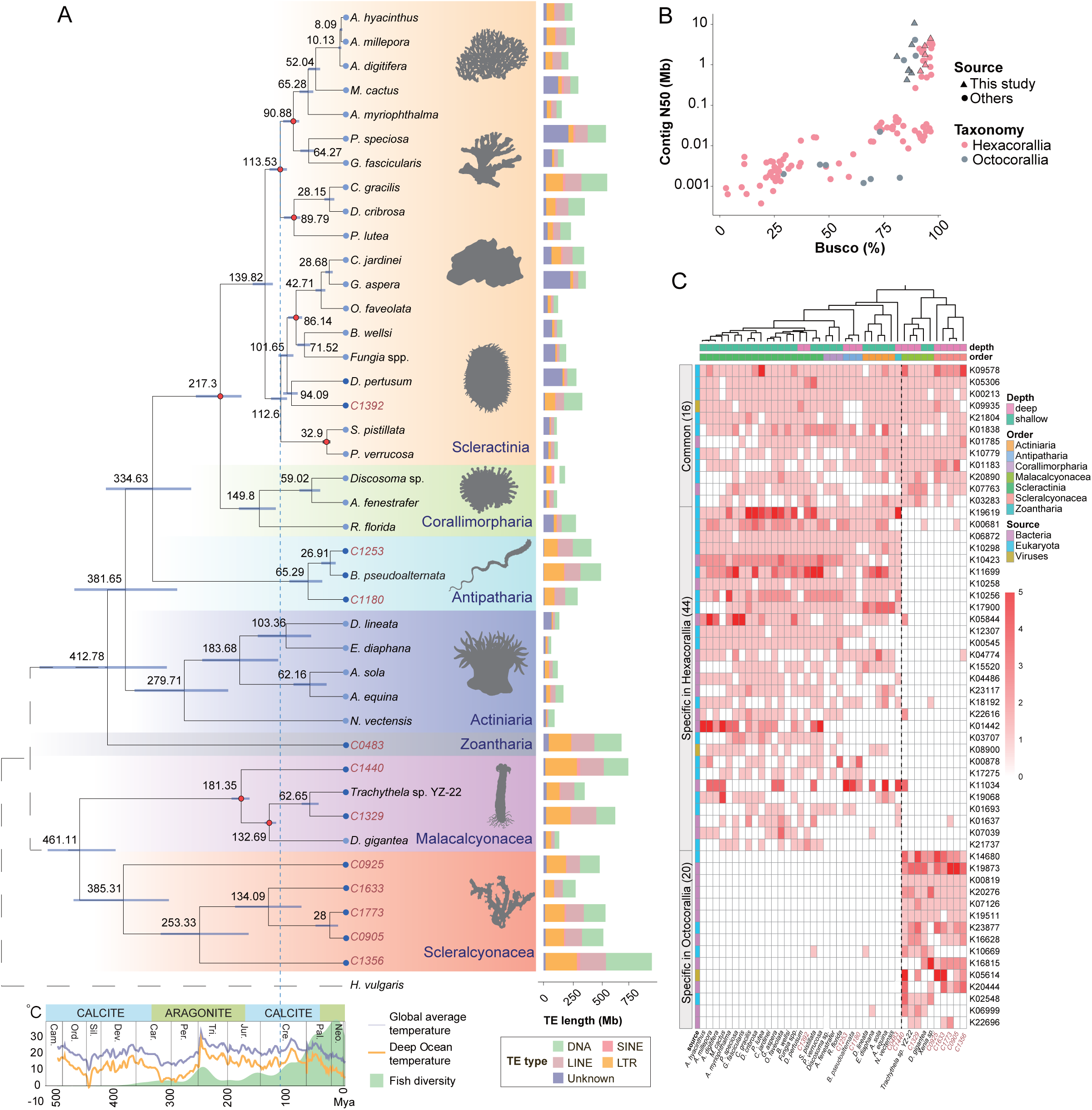
The evolution of coral species. (A) Phylogenetic tree of 41 coral species. Highlighted background colors denote different coral orders. The divergence time (MYA) of corals is labeled near nodes. Information about Earth’s temperature and fish diversity are shown on the timescale. The bar with green and blue indicates aragonite and calcite seas, respectively. The dotted line marks the speciation time of Scleractinia. The bars show the TE content (length in Mb) for each species. (B) The quality of assembled coral genomes was compared by contig N50 (y-axis) and BUSCO completeness score (x-axis). The 11 deep-sea corals sequenced in this study are shown as triangles. (C) Distribution of horizontal gene transfer (HGT) genes in coral species. Habitat type (shallow or deep) and orders are shown on the top, and the source of HGTs is shown on the left.

In the 11 deep-sea corals, 23,683–44,995 protein-coding genes were predicted, and gene statistics (including average gene length, exon length, and intron length) were found to be similar to those of shallow-sea corals (Table S4). In contrast, the 11 deep-sea corals show considerable variation in their repetitive sequences, with the lowest transposable elements (TEs) content at 34.13% and the highest at 63.77% (Table S5, Figure 1A and Figure S1A). Compared with other coral genomes, we found that Hexacorallia species have a shorter TE length than Octocorallia species, especially for LINEs, DNA transposons and LTRs (LINEs, average_Hexacorallia_ = 59.47 Mb, average_Octocorallia_ = 128.82 Mb; DNA transposons, average_Hexacorallia_ = 87.02 Mb, average_Octocorallia_ = 159.11 Mb; LTR, average_Hexacorallia_ = 54.02 Mb, average_Octocorallia_ = 150.02 Mb). We also observed that the proportion of repetitive sequences is significantly related to the genome size of corals (Figure S1A).

Taxonomic status was inferred through the integration of whole-genome sequencing data and cytochrome c oxidase subunit I (COI) gene phylogenetics (Figure 1A, Figure S2A). These species include one Scleractinia (C1392), two Antipatharia (C1253 and C1180) and one Zoantharia (C0483) from Hexacorallia, as well as two Malacalcyonacea (C1440 and C1329) and five Scleralcyonacea (C0925, C1633, C0905, C1773, and C1356) from Octocorallia (Figure 1A). Notably, phylogenetic analysis revealed moderate support from the COI gene tree for the placement of species C0483 within the Malacalcyonacea clade. In contrast, robust concordant support was obtained from whole-genome analysis and analysis of 1022 single-copy orthologous genes, both consistently resolving this species as an independent lineage and belong to Hexacorallia (Figure 1A, Figure S2B). Further morphological examination(Figure S2C-F) identified characteristic features of the order Zoantharia in both the skeletal cross-section and polyp structure^41,42^.Furthermore, we also provide morphological evidence for the remaining 10 coral individuals(Figure S3- S12). Among them, corals C1773 and C0905 exhibit similar morphology but inhabit water depths of 1773 m and 905 m, respectively.

### Co-evolution between corals and coral reef species and their holobionts

Coral reefs serve as both a habitat and source of food for coral reef species such as fish. To investigate the relationship between the timing of coral speciation and the number of reef species, we constructed a phylogenetic tree using 41 coral genomes from five Hexacorallia orders (Actiniaria, Antipatharia, Corallimorpharia, Scleractinia, and Zoantharia), and two Octocorallia orders (Malacalcyonacea and Scleralcyonacea). The phylogenetic analysis revealed that Hexacorallia species evolved towards greater diversity in response to changes in Earth’s environment and climate, while most Octocorallia species remained in deep-sea habitats (Figure 1A). Scleractinia, the most recently differentiated land-adapted and reef-forming corals, appeared about 217 million years ago in an ocean environment dominated by aragonite and high-Mg calcite precipitation and proliferated around ∼110 million years ago (MYA) (Figure 1A). Coincidentally, from the Late Cretaceous period, the number of fish species also dramatically increased (Figure 1A). This result suggests that during the Cretaceous period (145-66 MYA), the expansion of Scleractinia species increased coral reef species diversity, coinciding with rising global temperatures. Following the Cretaceous-Paleogene extinction event (∼66 MYA), coral and coral reef species underwent rapid proliferation and established their current distribution patterns (Figure 1A). The post-extinction period provided favorable conditions for coral diversification. Subsequently, corals provided the accessibility of space and food necessary for the proliferation of coral reef species. Demographic history analysis revealed a continuous decline in the effective population size of deep and offshore corals since 2 MYA (Figure S1B), which may be associated with changes in Earth’s temperature.

We next investigated the co-evolution of coral holobionts and detected 126 HGT events in corals from 41 phylogenetic clades (Figure 1C and Table S6). These HGTs mainly originated from bacteria (74 genes), fungi (16 genes), Alveolata (eight genes, including from Symbiodiniaceae), and Streptophyta (six genes) (Figure S1C). Among these, 16 HGTs are widely distributed in Hexacorallia and Octocorallia, 44 HGTs were only found in Hexacorallia, and 20 HGTs were only present in Octocorallia (Figure 1C). Of the 16 shared HGTs, four genes were transferred from Symbiodiniaceae to the coral ancestor and have been conserved in both clades (Table S6). This suggests that the common ancestor of Hexacorallia and Octocorallia had a symbiotic relationship with the ancestor of Symbiodiniaceae. Considering the reliance of Symbiodiniaceae on photosynthesis, the common ancestor of corals may have experienced an extended period of symbiosis in shallow seas, and some of them drifted to the deep seas during the ecological evolution of the earth and formed a new lineage. Based on genome data from a comprehensive lineages of deep-sea and shallow-sea corals at the order level, this study suggests that the history of symbiosis between coral and symbiotic algae may be much longer than the time of differentiation between Hexacorallia and Octocorallia.

### Symbiotic metabolic interactions of coral holobionts

To understand the nutrient recycling and retention of coral holobionts, we investigated the 2-Aminoethylphosphonate (AEP) and its N-alkylated derivatives widely present and abundant in natural phosphonates^43^. Microorganisms have been found to possess specific phosphate degradation systems for AEP, such as the AEP transaminase (*PhnW*) – Phosphonatase – *PhnX* system in coral reefs^44,45^. This system involves an initial transamination catalyzed by *PhnW*, the conversion of AEP into phosphonoacetaldehyde (PAA), followed by the conversion of PAA into acetaldehyde and inorganic phosphate through the action of the hydrolase *PhnX*^46^.

Interestingly, we found that the genes *PhnX* and *PhnW* are present in almost all examined coral genomes. Phylogenetic analysis showed that *PhnX* is transferred from a microbe to Symbiodiniaceae and, subsequently, incorporated into coral genomes (Figure 2A and Figure S13A). The collinearity results showed that *PhnX* is conserved in most coral genomes, indicating that this HGT event occurred on the common ancestral genome of corals (Figure S13B). Transcriptome data analysis of ten coral species showed that these two genes are expressed, indicating that corals can degrade AEP into acetaldehyde or inorganic phosphorus (P). The degradation of phosphate is particularly beneficial for organisms inhabiting marine environments where the bioavailability of phosphorus is often limited^47,48^. We propose that corals can metabolize dissolved organophosphorus, influencing the growth and composition of microbial communities in coral reef ecosystems and impacting the ocean’s nitrogen fixation and CO_2_ absorption capacity.

**Figure 2.**
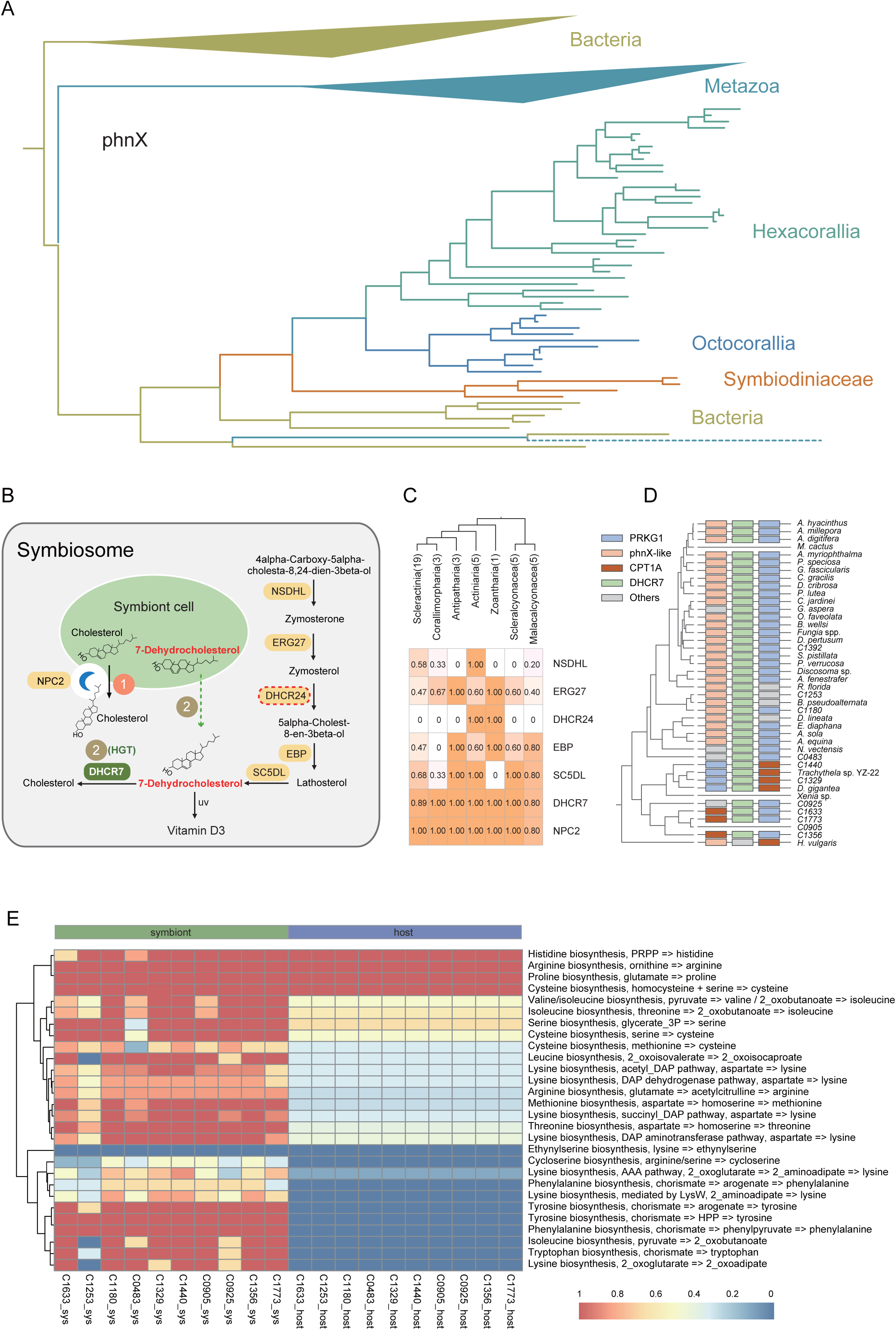
The co-evolution and nutrient utilization in different coral symbiotic community. (A) Maximum likelihood tree of PhnX. The coral clade is clustered with the Symbiodinaceae clade and separated from the other Metazoa. (B) A coral holobiont cell may have two ways to obtain cholesterol. One is transporting cholesterol synthesized by symbiotic cells through the NPC2 protein; the other is the horizontal transferred gene product DHCR7 in the coral holobiont cell. Protein names are on the left of the arrow. (C) Proportion of species in each order in which cholesterol synthesis genes are present. (D) Overview of the horizontal transferred gene DHCR7 and its flanking genes across coral species. (E) Amino acid biosynthesis genes in deep-sea corals and its symbiosis.

The cellular supply of cholesterol can be traced back to two sources: it is produced through synthesis or acquired through importation^49,50^. Previous studies have shown that in the cnidarian–dinoflagellate symbiosis, coral hosts rely on their algal symbionts to supply sterols. Similarly, we also found that the key enzyme Delta24-sterol reductase (*DHCR24*, EC:1.3.1.72) involved in cholesterol synthesis is lost in all Hexacorallia and Octocorallia species except for five Actiniaria and one Zoantharia species in the basal Hexacorallia clade (Figure 2B and C). These results indicate that *DHCR24* is generally lost in corals, leading to an inability of corals to synthesize cholesterol. Coral species rely on symbiotic microorganisms to acquire this essential compound by Niemann-Pick Type C2 (*NPC2*) sterol transporter (Figure 2B). However, we discovered a horizontal transferred gene from fungi, 7-dehydrocholesterol reductase (*DHCR7*) (Figure S13C). DHCR7 catalyzes 7-Dehydrocholesterol (7-DHC) to cholesterol in the final step of cholesterol synthesis^51^. Collinearity results showed that *DHCR7* is present in most coral genomes, indicating that this HGT event occurred in an ancestral species (Figure 2D). Since 7-DHC can move between cells^52^, we propose that 7-DHC produced by the endosymbiont can be transported into the host cell. Overall, our findings suggest that in addition to sterol transfer mediated by symbiont NPC2 proteins, corals use a pathway where 7-DHC is synthesized into cholesterol through the horizontal transferred gene *DHCR7* (Figure 2B).

In addition to investigating AEP and cholesterol metabolism, we also explored the amino acid synthesis pathways in deep-sea corals. To assess the completeness of amino acid biosynthesis across various deep-sea coral species, we constructed a comprehensive reference database, comprising 28 modules, 195 KEGG Ortholog (KO) entries, and 319,486 genes involved in amino acid synthetic pathways. Through this analysis, we identified 13,141 genes related to the biosynthesis of 17 essential amino acids within the genomes of deep-sea corals. Our findings revealed that 15 modules corresponding to ten amino acids (valine, tyrosine, tryptophan, threonine, serine, phenylalanine, methionine, lysine, leucine, and isoleucine) were fully present in the symbiotic microorganisms, but absent in the coral hosts. In contrast, the biosynthesis of proline, histidine, cysteine, and arginine was observed in both the host species and their symbiotic microorganisms (Figure 2E).

### The evolution of corals rhythm genes

Corals exhibit rhythmic phenomena such as calcification, tentacle expansion-contraction, and reproduction^53,54^. They retract their tentacles during the day and extend them to feed at night. For example, the feeding activity of the deep-sea Octocorallia *Paragorgia arborea* is highly seasonal, indicating that its biological rhythm is influenced by lunar cycles^55^. At the molecular level, the circadian clock system and biological rhythm are regulated through a conserved negative transcription-translation feedback loop controlled by five key genes^56^. Cryptochrome (*cry*) transcripts in corals show differential expression between day and night, suggesting a role in regulating simultaneous spawning behavior^57^. Another important gene involved in circadian rhythm regulation is *clock* (circadian locomotor output cycles kaput) and its homologous protein, neuronal PAS domain-containing protein 2 (*npas2*). Generally, *clock*/*npas2* and muscle ARNT-like 1 (*bmal1*) proteins act as positive regulators of endogenous circadian rhythms^58^, while *period* (PER) and *timeless* (TIM) genes serve as negative regulatory factors.

We comprehensively analyzed the fundamental rhythm genes in the genome of 41 coral species. The genes *bmal1* and *timeless* displayed conserved phylogenetic relationships and gene copies throughout coral evolution (Figure S14A and B). However, we noted the absence of *bmal1* in the deep-sea corals *C0483*, *C0905*, *C1773*, and *C1356* (Figure 3A), suggesting the independent gene loss events in these deep-sea corals. We also observed absence of *clock/npas2* and *cry* in deep-sea corals – four from Hexacorallia (*Bathypathes pseudoalternata*, *C1253*, *C1180* and *C0483*) and four from Octocorallia (*C0905*, *C0905*, *C1633* and *C1356*) (Figure 3A).

**Figure 3.**
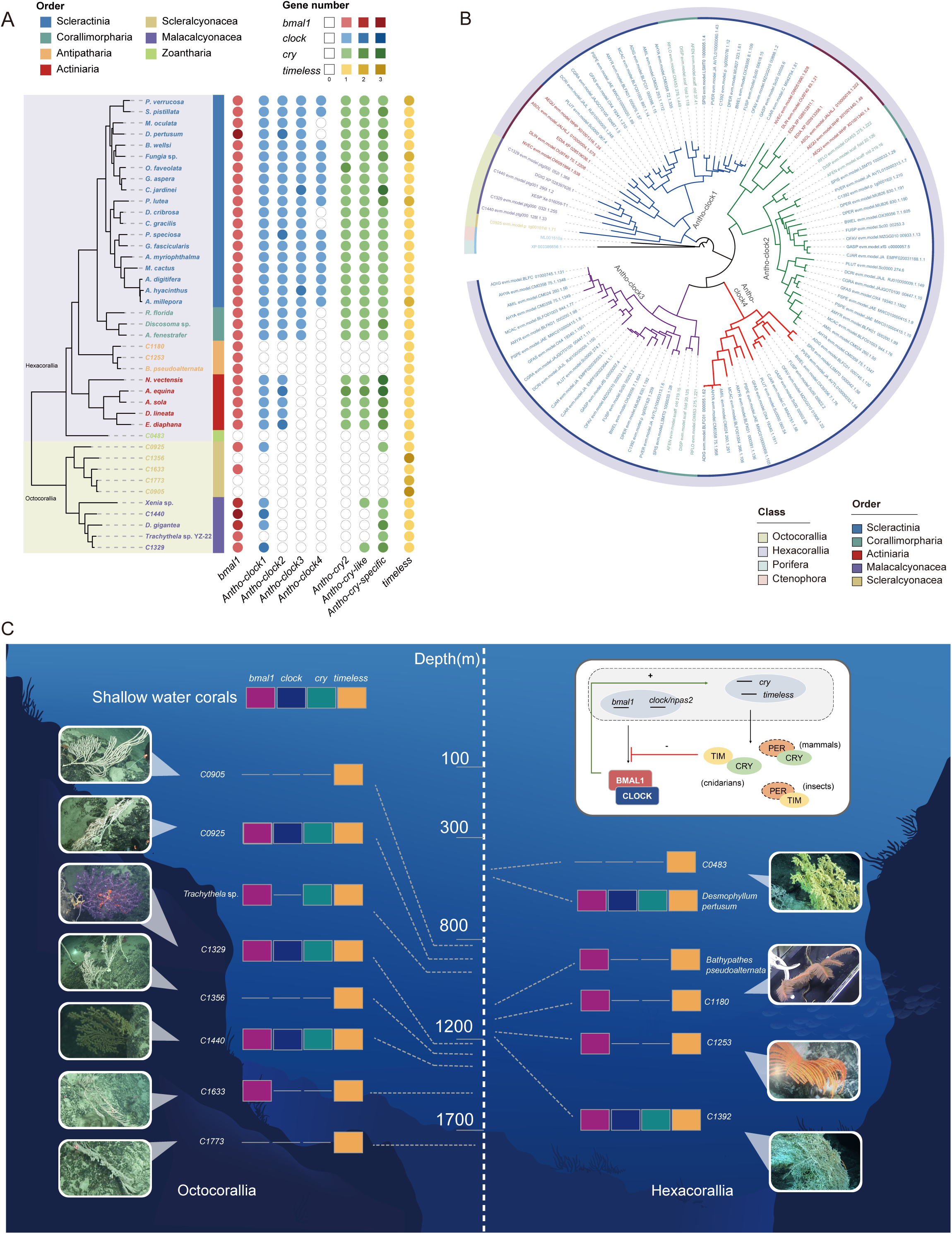
The distribution and evolution of key biological rhythm genes in coral genomes. (A) The gene numbers of *bmal1*, *clock*, *cry*, and *timeless* in coral genomes. Coral taxonomy of Hexacorallia (Scleractinia, Corallimorpharia, Antipatharia Actiniaria, Zoantharia) and Octocorallia (Malacalcyonacea and Scleralcyonacea) are indicated. (B) The phylogenetic tree of coral *clock* genes. From the outside to the inside, different colors represent the class and order of coral species, as well as four subtypes of the *clock* gene. (C) Combination pattern of four core rhythm genes in deep-sea corals sequenced in this study. The depth information and photos for each coral species are shown. A schematic diagram of the circadian clock BMAL1-CLOCK heterodimer and its transcription and translation feedback loops are shown in the white box.

We classified the *clock* genes in Anthozoan into four subfamilies: *Antho-clock1*, *Antho-clock2*, *Antho-clock3*, and *Antho-clock4* (Figure 3B). We observed four copies of *clock* genes in coral genomes, except for specific gene loss events in some deep-sea corals, in contrast to only one *clock* gene copy in the outgroup species *Amphimedon queenslandica* and *Mnemiopsis leidyi* (Figure 3B). The *Antho-clock1* subfamily represents the ancestral gene found in all corals. A presumptive gene loss event of *Antho-clock2* occurred at the divergence of the Octocorallia. The *Antho-clock3* gene is specific to Scleractinia and Corallimorpharia, while *Antho-clock4* is only present in Scleractinia (Figure 3B). However, we observed an exception where one copy of the *Antho-clock4* gene was lost in the deep-sea corals *C1392* and *Desmophyllum pertusum* in Scleractinia (Figure 3A) supporting the notion that these species retained certain genetic remnants of their shallow-sea ancestors during their evolutionary transition back to deep-sea habitats. Collectively, these results demonstrate the complex evolutionary dynamics of *clock* genes in coral species and highlight the influence of environmental adaptation on the presence or absence of these genes.

We also classified the cryptochrome (*cry*) genes of corals, insects, and mammals into three subfamilies: *Antho-cry-specific*, *Antho-cry-like*, and *Antho-cry2* (Figure S14C). We identified the *Antho-cry-specific* subfamily in Hexacorallia and Octocorallia corals as the ancestral gene copy of the *cry* gene in Anthozoa (Figure 3A). Interestingly, we only detected *Antho-cry2* in Scleractinia and Actiniaria, corals that primarily inhabit shallow-sea habitats, while the gene was absent in deep-sea corals (Figure S14C). *Cry-like* is a light-independent suppressor in the negative feedback loop in rhythm regulation, while *cry2* acts as an entrainment oscillator during light/dark transitions^59^. Based on the phylogenetic relationship, we speculate that during the evolutionary process, most deep-sea corals in Antipatharia, Zoantharia, and Scleralcyonacea did not gain *Antho-cry2* genes, except for *C1392* and *Desmophyllum pertusum*, which possibly were found in a shallow-sea habitat during their evolution. In corals, we did not find any *per* genes, critical regulators of clock gene transcription that interact with *cry* and *timeless* proteins. Only *cry* and *timeless* genes were found, suggesting that the negative feedback mechanism of the circadian clock of corals differs from that of mammals and insects (Figure 3C).

We observed that the presence and absence patterns of biological rhythm genes in shallow-sea corals tended to be consistent, while deep-sea corals present diverse patterns (Figure 3C and Figure S14D). These results highlight the complex evolutionary dynamics of biological rhythm genes in coral genomes, with variations in gene presence, loss or absence caused by genome rearrangement or pseudogenization occurring across different coral species, especially in deep-sea lineages (Figure S14E and F).

### Coral temperature adaptations

Shallow-sea corals are particularly susceptible to temperature changes, while deep-sea corals reside in cooler environments with relatively stable temperatures (around 4 ℃).Since Hexacorallia has the largest number of genomes and contains reef-building corals, and the bleaching of reef-building corals caused by rising temperatures is an ecological issue of great concern, we focused on Hexacorallia species to understand the genetic basis of temperature adaptation. We identified 180 genes with copy number variations between deep-sea and shallow-sea corals (Figure 4A). We further investigated the expression patterns of 117 genes with more copies in the shallow-sea coral genomes of Hexacorallia using published *Galaxea fascicularis* transcriptome data^60^. Copy number reduction of these genes during the transfer of some corals from shallow to deep-sea indicates that these genes may be critical for corals to adapt to shallow-sea temperatures. Interestingly, we found that 16 genes with more copies in shallow-sea corals showed higher expression levels in response to thermal challenges (Figure S15A), including the heat shock protein 70 kDa (*Hsp70*) and small heat shock/ α-Crystallin (*Cryab*) (Figure 4B). During heat stress, *Hsp70*, as a highly conserved molecular chaperone, plays a crucial role in facilitating the folding process of both newly synthesized proteins and those that have become denatured^61^. In addition, high expression of *Cryab* has been reported to have the ability to inhibit irreversible denaturation of other proteins and serve as an initial defense against heat stress^62,63^.

**Figure 4.**
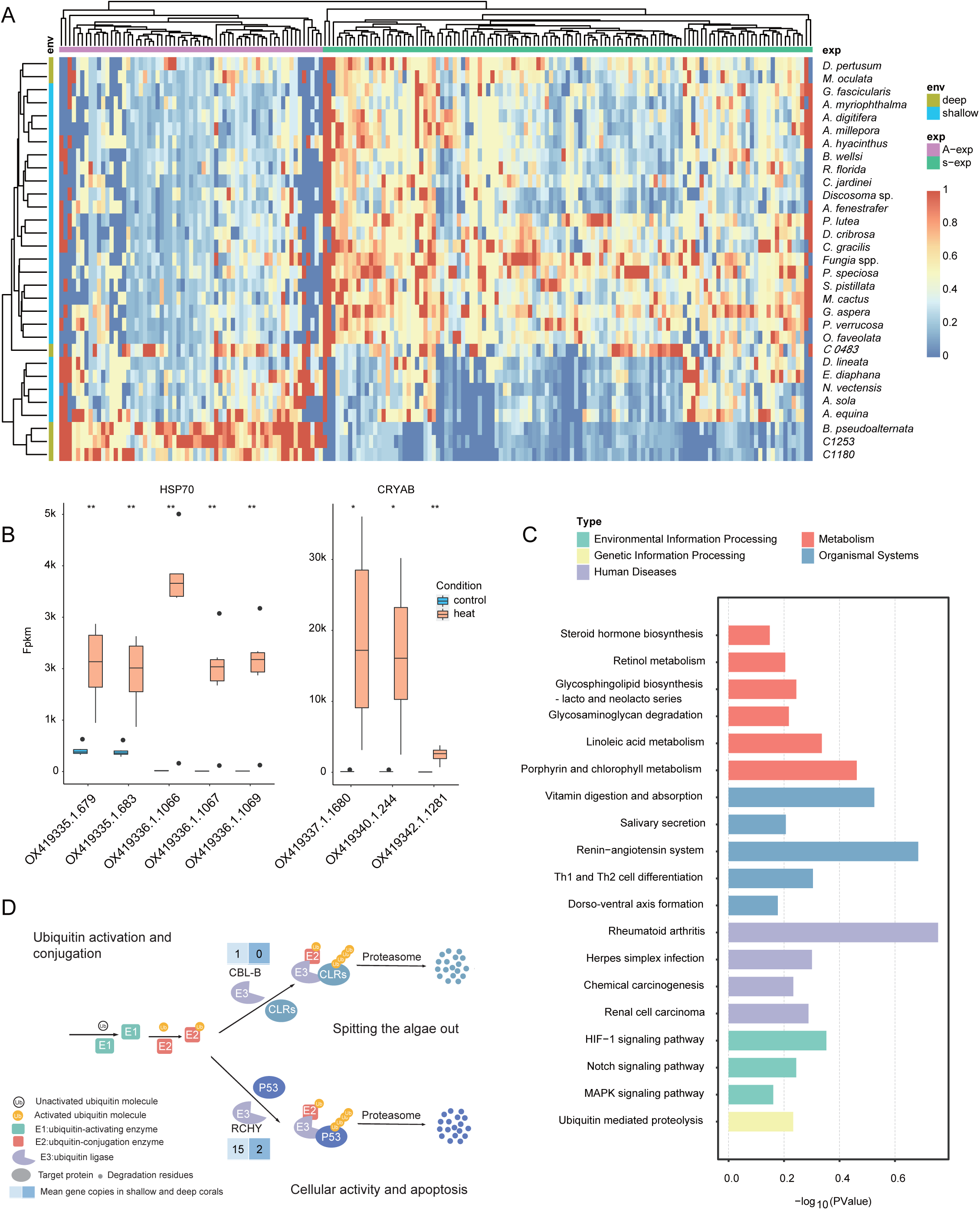
Genes related to temperature response in coral genomes. (A) Copy number distribution of genes with different expression patterns in response to temperature changes in deep-sea and shallow-sea corals. (B) The expression of the HSP70 and CRYAB genes of Galaxea fascicularis under control and heat conditions. (C) Gene enrichment of genes differently expressed in a Galaxea fascicularis temperature response experiment. (D) General view of ubiquitin-mediated proteolysis. The average gene number of E3 ubiquitin ligase Rchy and Cbl-b genes in shallow and deep coral genomes is shown.

We also performed functional enrichment of the above 180 genes, revealing enrichment for vitamin digestion and absorption, steroid hormone biosynthesis, the Notch signaling pathway, and ubiquitin-mediated proteolysis pathways (Figure 4C). Of particular interest, the ubiquitination process can increase protein degradation pathways to reduce temperature sensing^64^. Ubiquitination is known to mark damaged or misfolded proteins for degradation^65^. Heat stress causes protein misfolding, triggers cellular responses, and significantly increases protein ubiquitination^66^. Ubiquitination is a multi-step process that includes activation by E1 ligase, conjugation by E2 ligase, and conjugation to substrate proteins through E3 ligase. Interestingly, we observed notable differences in the gene copy numbers of *Cbl-b* (E3 ubiquitin-protein ligase) and *Rchy* (RING finger and CHY zinc finger domain-containing protein) within the ubiquitin-mediated proteolysis pathway between deep-sea corals and shallow-sea corals (Figure 4A). Specifically, the gene *Cbl-b* is absent in deep-sea corals but present in almost all shallow-sea corals (Figure 4D). CBL-B is known to activate the ubiquitination of C-type lectin receptors (CLR), which negatively regulates CLR and can trigger pro-inflammatory responses to microbial infection^67^. We propose that the presence of CBL-B in shallow-sea corals may be related to the coral bleaching process, where corals lose their Symbiodiniaceae due to heat stress. In addition, the gene copy number of *Rchy* in shallow-sea corals is seven-fold higher than in deep-sea corals (Figure 4D). This gene targets the degradation of the critical tumor suppressor and transcription factor p53, and recent studies on mammalian cells and mice highlight a relationship between p53 and temperature response^68-70^. Within the temperature threshold, p53 is thought to promote cell survival by suppressing excessive heat shock responses, but above a threshold temperature, it exacerbates apoptosis of damaged cells^68^. Our results suggest that the *Cbl-b* and *Rchy* allow shallow-sea corals to adapt to the heat stress caused by global warming.

## Discussion

In this study, we collected, sequenced, assembled, and annotated the genomes of 11 deep-sea corals. These included four Hexacorallia, seven Octocorallia species, and three orders (Antipatharia, Zoantharia, and Scleralcyonacea) with significant genomic gaps. These assemblies greatly expand the genome resources of deep-sea corals.

Although deep-sea corals live in dark, low-oxygen, and low-temperature environments, their genetic diversity is very high. It was difficult to detect the genetic characteristics of deep-sea corals and shallow-sea corals from the genome data. Most gene family differences result from phylogenetic differences rather than sea-level habitats. Subsequently, we compared the genomic differences between deep-sea and shallow-sea corals in Hexacorallia and Octocorallia corals. In Octocorallia, the expanded gene families in the deep sea showed enrichment for gene ontology terms such as “protein phosphorylation and protein metabolic process” (BP), “voltage-gated potassium channel complex” (CC), and “protein kinase activity” (MF) (Figure S15B). In Hexacorallia, the expanded gene families were enriched for the “transforming growth factor beta receptor signaling pathway” (BP), “membrane” (CC), and “hydrolase activity, acting on glycosyl bonds” (MF) (Figure S15C). These data indicate that Hexacorallia and Octocorallia corals evolved distinct strategies to adapt to a deep-sea environment.

Our study also provides valuable insights into the relationship between coral reef species and the timing of coral speciation. Coral reefs are important habitats and food sources for a diverse range of species. Our study shows how they correlate with species radiation and evolutionary diversification. By analyzing the genomes of 41 different coral species, we found that Hexacorallia species evolved towards greater diversity in response to changes in Earth’s environment and climate, while most Octocorallia corals remained in the deep sea. The emergence of Scleractinia, the most recently differentiated land-adapted and reef-forming corals, might drive coral reef species diversity. We highlight the importance of understanding the historical dynamics of coral speciation and the influence of environmental factors on the evolution of coral reef ecosystems. By emphasizing conservation efforts to protect these ecosystems, this study has important implications for the future of coral reefs and the species that depend on them. While coral evolution involves complex genomic events such as gene duplication, horizontal gene transfer (HGT), and recombination. Whole-genome data are more likely to detect these events, thus revealing evolutionary trajectories distinct from those inferred from mitochondrial markers or ultraconserved elements (UCEs)^29^. Future studies should integrate larger-scale genomic datasets from both deep-sea and shallow-water corals, along with more sophisticated analytical methods, to resolve these discrepancies and provide a more accurate understanding of coral evolutionary history.

HGT events are important for the evolution and adaptability of organisms because they facilitate adaptation and the acquisition of new genetic information from different species^71^. We found the horizontal transferred gene *PhnX* in the coral genomes, suggesting the presence of an AEP-specific phosphate degradation system in corals^44,45^. This system is particularly beneficial for organisms living in marine environments where phosphorus availability is often limited^47,48^. These results highlight the importance of HGT in shaping coral adaptation and survival strategies. Moreover, four genes transferred from Symbiodiniaceae to the coral ancestor and preserved in both clades strongly suggest that the common ancestor of Hexacorallia and Octocorallia had an established symbiotic relationship with Symbiodiniaceae. Given the reliance of Symbiodiniaceae on photosynthesis, it is highly likely that ancestral corals experienced a prolonged period of symbiosis in shallow seas, and some corals eventually evolved to be a distinct lineage after drifting into deep seas. Employing genomic data deep-sea and shallow-sea corals at the order level, our study demonstrates that the symbiosis history of corals and symbiotic algae is much longer than previously known.

The symbiosis between corals and algae or microorganisms has been widely studied. We here focused on metabolic complementary processes between corals and their symbionts. Corals rely on algal symbionts to provide sterols rather than synthesizing them themselves^24^. We found that corals may have lost the Delta24-sterol reductase gene, resulting in an apparent inability to synthesize cholesterol by the coral species. However, our study suggests that the horizontal transferred gene *DHCR7* allows the uptake and formation of cholesterol from 7-DHC produced by the endosymbiont. This finding reveals a pathway for corals to obtain cholesterol in addition to uptake from symbionts. Amino acids provided by symbiotic algae have been reported in shallow-sea corals^72^. We reveal a similar scenario in deep-sea corals. Some amino acids in deep-sea corals are likely obtained through their symbiotic bacteria. The metabolic complementarity between these symbionts and the coral host has great biological and ecological significance for coral research.

Our analysis of biological rhythm genes in coral genomes revealed complex evolutionary dynamics in coral species manifested as differential gene presence, duplication, and absence across species and lineages. The genes *bmal1* and *timeless* show a relatively conserved pattern in their presence and evolutionary relationships in corals. The *clock/npas2* gene exhibits quantitative differences between shallow- and deep-sea corals, suggesting habitat-specific patterns. Phylogenetic analysis revealed unique *clock* gene subtypes in stony corals, suggesting the presence of gene duplication in their common ancestor. The absence/presence pattern and presentation (gene copy number) of *cry* genes in deep-sea corals indicate significant changes and adaptations in response to environmental stressors.

Understanding the genetic mechanisms behind temperature adaptations of corals remains critical for their protection. By analyzing gene copy number variations and differential gene expression in response to temperature changes, we identified *hsp70* and *Cryab* as candidate coral temperature adaptation genes. We also found copy number differences of the genes *Cbl-b* (E3 ubiquitin-protein ligase) and *Rchy* (RING finger and CHY zinc finger domain-containing protein) between deep-sea corals and shallow-sea corals. These genes may participate in the thermal stress process of corals through the ubiquitin-mediated proteolysis pathway. Further research is needed to fully comprehend the functional significance of these genes and their potential for improving coral conservation efforts in the face of climate change.

Overall, this research expands our knowledge of corals from multiple aspects, including evolution, symbiosis, and environmental adaptation, by sequencing deep - sea coral genomes and conducting comparative genomics analysis. It provides crucial insights for understanding coral diversity and their potential to cope with environmental changes.

## Materials and Methods

### Sample collection

From December 2020 to June 2022, deep-sea corals were collected from a cold-water coral forest in the South China Sea (SCS) using the human-occupied vehicle (HOV) Fendouzhe and Shenhai Yongshi. The corals were collected using a handnet and placed in a biobox. Immediately after the dives, the coral samples were washed with 1× PBS and transported to -80 ℃ for preservation.

#### Morphological identification

Skeleton and polyp of corals were soaked in sodium hypochlorite for 15 minutes, then centrifuged at 1 000 r/min for 2 minutes at room temperature. The resulting sediments were subsequently rinsed with distilled water and 70% ethanol to collect the sclerites. These sclerites were air-dried, placed on petri dish for Electron microscopic observation.

### DNA and RNA extraction and sequencing

To extract total DNA, small pieces of coral colonies were ground using a sterile mortar with liquid nitrogen. The CTAB method was used for DNA extraction, and a Genomic-tip 100/G kit (QIAGEN, USA) was used for purification. The quality and quantity of the DNA samples were checked using gel electrophoresis and a Qubit 2.0 Fluorometer (Life Technologies, USA), respectively. For whole genome sequencing (WGS) libraries, 1 ug of high-quality DNA was used to construct WGS libraries using an MGIEasy Universal DNA Library Prep Set (MGI, China). The WGS libraries were sequenced on the MGI2000 platform. Additionally, a 20-kb circular consensus sequencing (HiFi CCS) library was constructed using 15 ug of high-quality DNA from one individual, using the SMRTbell Express Template Prep Kit 2.0 (Pacific Biosciences, PN 101-853-100), and sequenced on a PacBio Sequel sequencer. To extract total RNA, a TRIzol kit (Invitrogen, USA) was used following the manufacturer’s protocol. The quality and quantity of the total RNA sample were evaluated using gel electrophoresis and Qubit 2.0 Fluorometer (Life Technologies, USA), respectively. An MGIEasy rRNA Depletion Kit (MGI, China) was used to capture the total mRNA from 1 ug of total RNA. The captured total mRNA and eukaryotic mRNA were then utilized to construct cDNA libraries using the MGIEasy RNA Library Prep Set (MGI, China) and sequenced on the MGI2000 platform.

### Genome assembly and evaluation

For the WGS data, SOAPnuke v1.6.5^73^ was used to filter the reads of low quality, adapter sequences or duplication with the parameters “-A 0.2 -M 2 -l 10 -q 0.1 -n 0.05 -Q 2 - d”. The obtained clean data were used for 21-mer analysis using jellyfish v2.2.6^74^.Then GenomeScope2^75^ was used for estimation of genome sizes and heterozygous for each coral species.

The PacBio HIFI data were assembled using hifiasm v0.14.1-r314^76^. bwa v0.7.12^77^ was used to map the WGS data back to the primary assembly, while purge_dups v1.2.5^78^ pipeline was used to remove redundancy sequences. The purged assembly was interrogated using BUSCO (Benchmarking Universal Single-Copy Orthologs) V5^79^ pipeline with the metazoa_odb10 database.

### Genome annotation

LTR_FINDER v1.0.6^80^ and RepeatModeler v1.0.8^81^ for de novo repeat sequence annotation. ProteinMask, TandemRepeatFinder v4.07^82^ and RepeatMasker v4.0.6^83^ were used homology-based repeat annotation. The gene models of the newly sequenced deep-sea genomes in this study were predicted by homolog, de novo, and transcriptome approaches. For homolog prediction, the protein sequences the corals *Acropora digitifera* (GCF_000222465.1), *Pocillopora damicornis* (GCF_003704095.1), and *Acropora millepora* (GCF_004143615.1) were selected as reference to compare with each of our 11 genomes and gene models were predicted using GeneWise v2.4.1^84^. For de novo prediction, Augustus v3.1.0^85^ and GeneMark v4 ^86^ were used to scan for candidate gene models. For transcriptome data, the RNA reads were mapped against the genome sequences using HISAT v2.1.0^87^ to align RNA sequencing reads back to the genome, then StringTie v1.3.5^88^ and TransDecoder v5.5.0^89^ were used to predict the expressed genes. Finally, EvidenceModeler^90^ is used to integrate the results of homology, de novo and transcriptome-based predictions. Next, we removed gene models (EVMs) with a 90% overlap with repeat sequences, only supported by the de novo approach without function, or with N bases of coding sequences larger than 30%. Finally, EVM amino acid sequences were compared against the metazoa_odb10 database using the BUSCO V5 pipeline protein mode to evaluate the quality of the gene set. Protein function was predicting alignments to KEGG, UniProt and NR databases using diamond v0.9.21.122^91^, and domain information were annotated using InterProScan v59-91^92^.

### Phylogenetic analysis

The genome phylogenetic trees were constructed using gene sequences from 40 species (the 11 novel genomes in this study and 29 published coral genomes). Firstly, OrthoFinder v2.3.8^93^ was used to identify homologous genes, generate gene families and determine approximate phylogenetic relationships among the whole species. Since the coral lineage has speciated for a long time a small number of shared single-copy genes than expected. We, therefore, extracted single-copy genes from the Hexacorallia and Octocorallia corals separately. Then we used MUSCLE v3.8.1551^94^ to construct multiple sequence alignment for all the single-copy gene families. The corresponding sequences from same species were concatenated, and trimal^95^ was used to filter those sites with gap ratio greater than 80%. Based on the retained alignments, IQ-TREE v2.1.4^96^ was applied to construct a maximum likelihood tree (parameter -m MFP -B 1000 --date-ci 200). Finally, MCMCTREE v4.9j^97^ is used to infer the divergence time of coral species with the reference fossil time that taken from TimeTree (parameters ndata=3 seqtype=0 clock=3 model=4 cleandata=0 burnin=2000 samfreq=2 nsample=20000).. The expansion and contraction of gene families were estimated using CAFE5^98^.

For the construction of the COI phylogenetic tree, all cytochrome c oxidase subunit I (COI) protein sequences belonging to the Anthozoa lineage from the NCBI mitochondrion protein sequence database were extracted. These sequences served as references for annotating this gene in all unfiltered coral genome assemblies using miniprot. The annotated genes were then aligned against the reference sequences using diamond and the gene with the highest alignment score against the reference was selected for each species. Multiple sequence alignment of all COI genes was performed using muscle with default parameters, followed by alignment trimming using trimAl with parameters -gt 0.8 -resoverlap 0.8 -seqoverlap 80. Subsequently, phylogenetic tree construction was conducted using FastTree, and the resulting tree was visualized using an R script.

To further clarify the phylogenetic position of C0483, we selected 1–2 representative species from each order within Anthozoa to construct a backbone phylogenetic tree. Hydra vulgaris was used as the outgroup. A total of 1,022 single-copy orthologous genes were identified across these species. Gene trees were inferred for each orthogroup using IQ-TREE, and a consensus species tree was subsequently reconstructed using ASTRAL-Pro 3.

### Identification of HGT events

Firstly, the amino acid sequences of 41 corals were aligned to the NR database (2022-11) using diamond v0.9.21.122 (--evalue 1e-5 --more-sensitive --masking none). Hits that come from Anthozoa are removed, and the best 50 hits for every genes were retained and their taxonomy classifications were count. Those genes that does not include any hits from Metazoan sequences were regarded as candidate HGTs. To eliminate potential contamination and low-quality sequences, the proportion of genes in the coral genome sequences with Metazoan best hit was calculated. If more than half of the genes on the genome sequence were with non-Metazoan best hits, or the sequence length was less than 10k, it is considered that this sequence is more likely to be contaminated or has low credibility, and the genes on this sequence were deleted from the candidate HGT list.

To minimize the impact of different gene prediction methods on the outcomes, coral genes potentially representing HGT were re-predicted using a homology-based prediction approach. Specifically, all remaining coral genes related to candidate HGTs and their top 50 NR hits were combined together, then self-aligned with diamond and clustered with MCL^99^ (parameter: -I 1.5). Orthogroups (OGs) that contained more than three sequences, including at least one coral gene were retained. These OGs were then deduplicated using CD-HIT ^100^ (-c 0.9 -G 1), and the resulting sequences were used as a non-redundant reference library. Based on this library, genes were predicted on the 41 coral genomes using miniport v0.11^101^, yielding new HGT candidate genes with OG tags. These genes were re-aligned to the NR database using Diamond (--max-target-seqs 0 --more-sensitive --masking none), and only best hit in each order was retained.

To determine the HGT origins, the NR sequences in each OG were categorized based on their phylogenetic information according to NCBI taxonomy database. Briefly, sequences belonging to the superkingdoms of Viruses, Bacteria, and Archaea are directly classified into the superkingdom to which they belong, and sequences belonging to the superkingdom of Eukaryota are classified into the secondary subcategories under Eukaryota to obtain a more detailed classification. Any category within an OG that had fewer than two sequences was discarded. MAFFT v7^102^ was used to perform multiple sequence alignment for each OG (--auto), and trimAl v1.4 was used to delete the sites with excessive gaps and sequences that with poor consistency (-- gt 0.5; -resoverlap 0.5 -seqoverlap 50). OGs that (1) with less than two coral genes, or (2) with less than 3 NR sequences, or (3) more than 30% of the sequences with more than 80% of the components are composed of less than 5 amino acids were removed. Phylogenetic trees were constructed for the remaining OGs using FastTree v2.1.11^103^, and then median rooting, visualization and collapse were performed using the R packages *ape*, *phangorn*, *ggtree*, *bio*, and *ete3*. The color of the branches was determined by their category, and the nodes with the same category were collapsed.

Finally, all the OGs’ phylogenetic trees were manually screened to determine whether genes were from HGT. Considering the incompleteness of the database, we mainly use the overall tree lineage structure to determine whether the coral gene is HGT or not, and used the class information of sister branches of the coral gene cluster to determine the source of HGT. This means that some genes that did not have a nested structure were also considered to be HGTs, and the source of few genes that were considered to be HGT could not be determined.

### Identification of amino acid pathway genes

Contigs with more than half of the genes annotated as microorganisms were removed to reconstruct the clean draft genomes of corals. The CCS data were mapped to the clean draft genomes using minimap2^104^ with default settings, and then the samtools^105^ was used to extract the un-host CCS reads. The genes involved in the synthetic pathways of eighteen amino acids were collected from the KEGG database. The un-host CCS reads were used to search the potential genes associated with amino acids synthetic against reference eighteen amino acids database using diamond with the parameters ‘--evalue 1e-5 -k 0 -p 10 --sensitive’. The genes that with more than 30% identity and 80% length coverage were screened as the target genes for further analysis. The in-house scripts were used to calculate the completeness of eighteen amino acids synthetic pathways.

### Rhythm gene analysis

This study mainly identifies and analyzes five important rhythm genes: *bmal1*, *clock*, *cry*, *period*, and *timeless*. First, we searched for the five target gene lists from the function annotation results and calculated the number of genes for each species. Since some corals do not have hits and all corals didn’t have *period* genes, we further downloaded the target genes NCBI NR database and combined them with the identified coral gene lists. The obtained 2472. 4563, 4562, 6408, and 3807 sequences for *bmal1*, *clock*, *cry*, *period*, and *timeless* genes were used as reference libraries, and miniprot was used to predict the homologous genes on all the 41 corals genomes. In this way the genes were supplemented and revised. The final gene lists were used to calculate and display the presence and distribution in all coral genomes. At the same time, the genes were aligned using MAFFT v7, and gene trees were constructed by FastTree.

## Data Availability

The genomic data that generated in this study are available at CNSA (https://db.cngb.org/cnsa/) under the accession CNP0004936.

The code for the custom scripts for data analysis is available from a github repository: https://github.com/BGI-Qingdao/Module_completeness

## Abbreviations

HGT: horizontal gene transfer
BUSCO: Benchmarking Universal Single-Copy Orthologs
TE: transposable element
LINE: Long interspersed nuclear element
LTR: Long terminal repeat
MYA: Million years ago
KO: KEGG Ortholog
SCS: South China Sea
HOV: Human-occupied vehicle
WGS: whole genome sequencing
CCS: circular consensus sequencing
OG: Orthogroup;

## Acknowledgement

This work was financially supported by the National Key R&D Program of China (2022YFC2805404), the Project of Sanya Yazhou Bay Science and Technology City (Grant No: SKJC-2024-01-002), and the Technological Innovation Projects of Qingdao West Coast New Area (ZDKC-2022-03). This study was also supported by the High-performance Computing Platform of YZBSTCACC (YaZhou Bay Science and Technology City Advanced Computing Center). Additionally, we are grateful to the captain, crew, and scientific staff of the R/V Tansuo 1 and Tansuo 2 and the pilots of the manned submersibles Fendouzhe and Shenhaiyongshi. We are also grateful to the China National GeneBank for the data production.

