## Supplemental Data 1 for "Eleven deep-sea coral genome assemblies unveil insights into evolution, adaptation, and coral biodiversity"

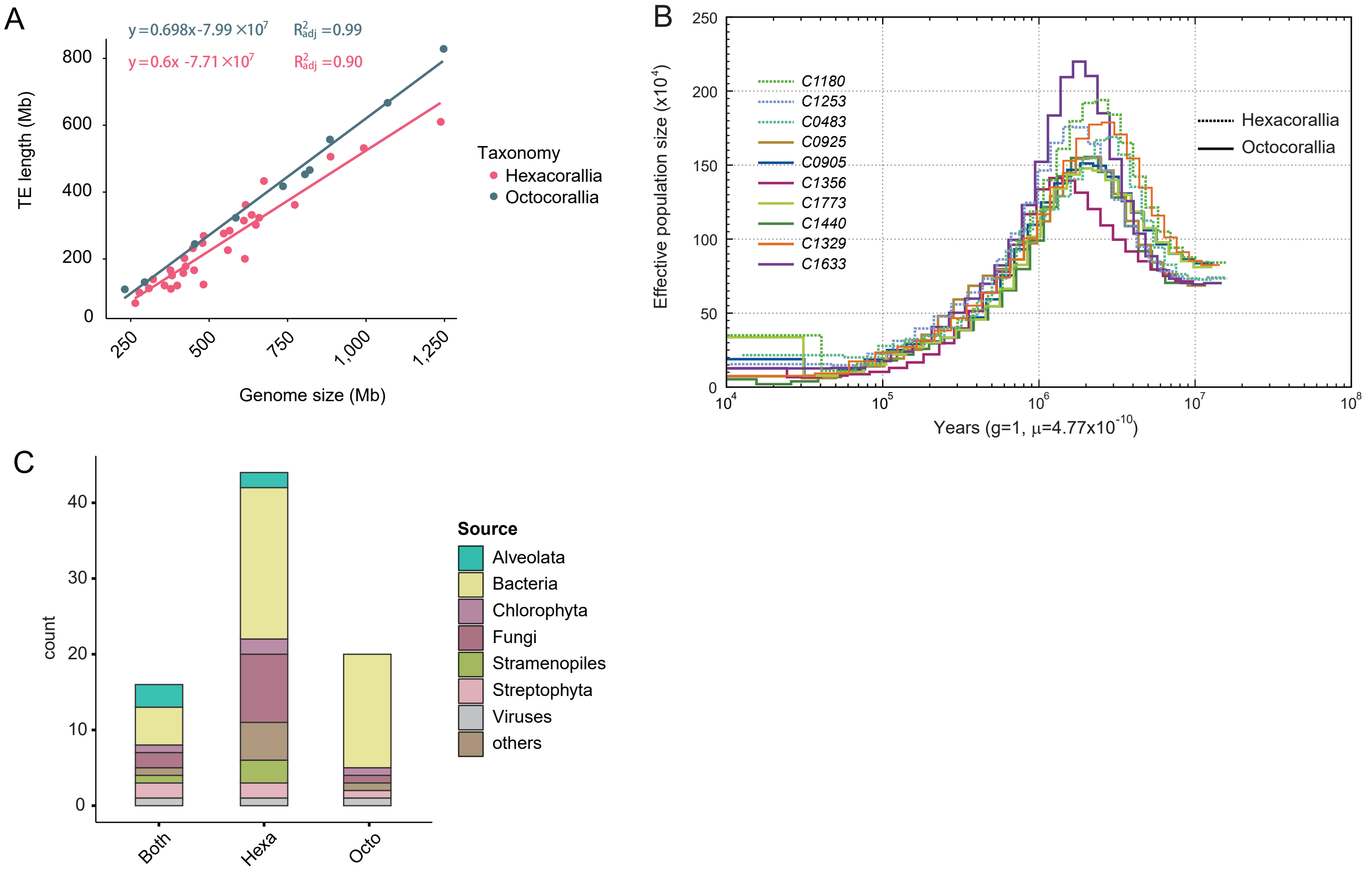


**Fig. S1.** Statistics on characteristics of coral genomes and horizontal transferred genes. (A) Comparison of genome size and transposon content among 41 corals. (B) Demographic history of deep corals inferred using PSMC. (H) The origin taxonomy of 80 HGTs widely distributed in corals. (C) Distribution of horizontal transferred genes'functions.

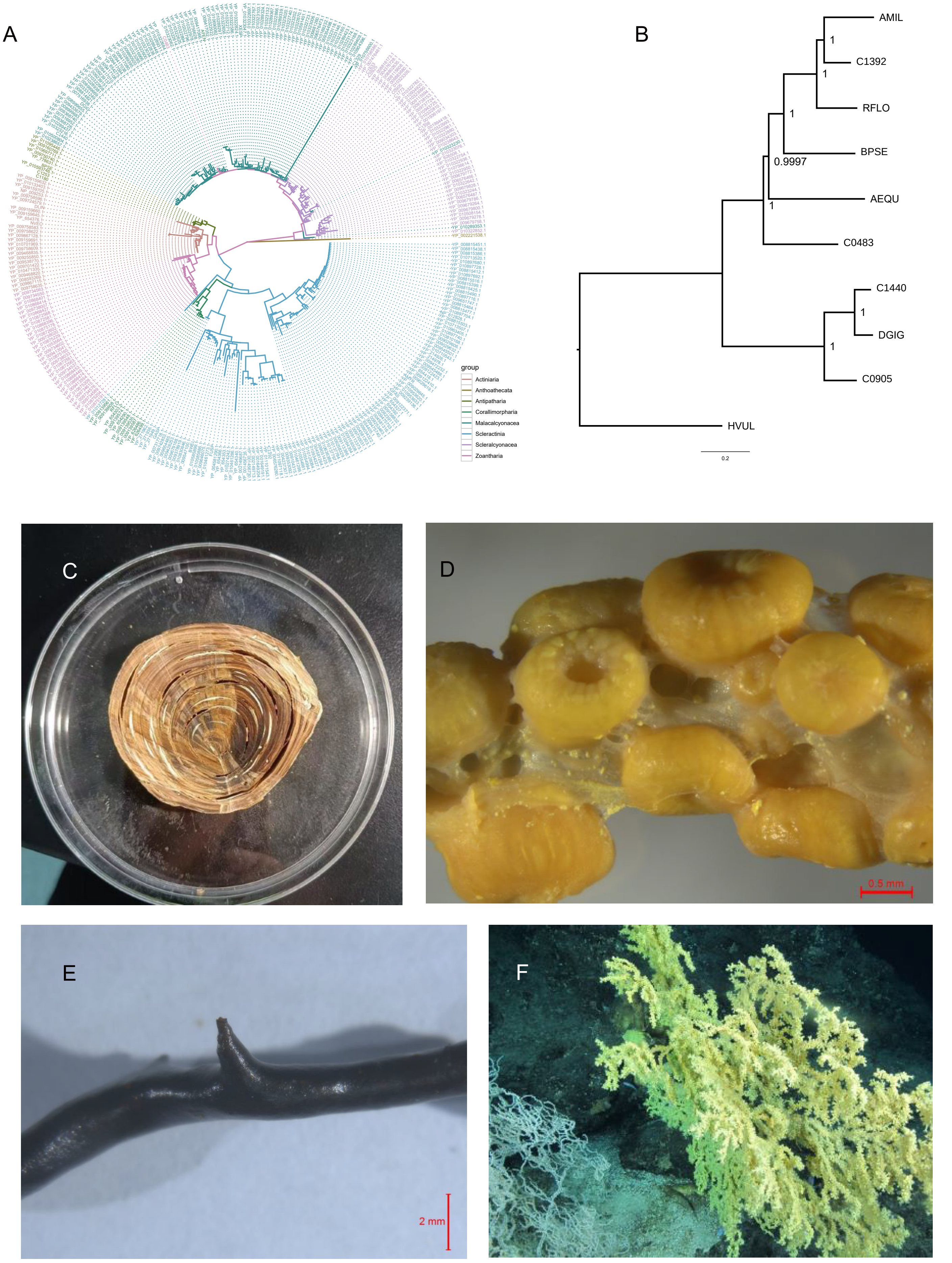


**Fig. S2.** Further identification of C0483.(A) Phylogenetic tree of Anthozoa COI genes. (B) Backbone tree of Anthozoa constructed using 1,022 single-copy genes. (C-F) Morphology of C0483.

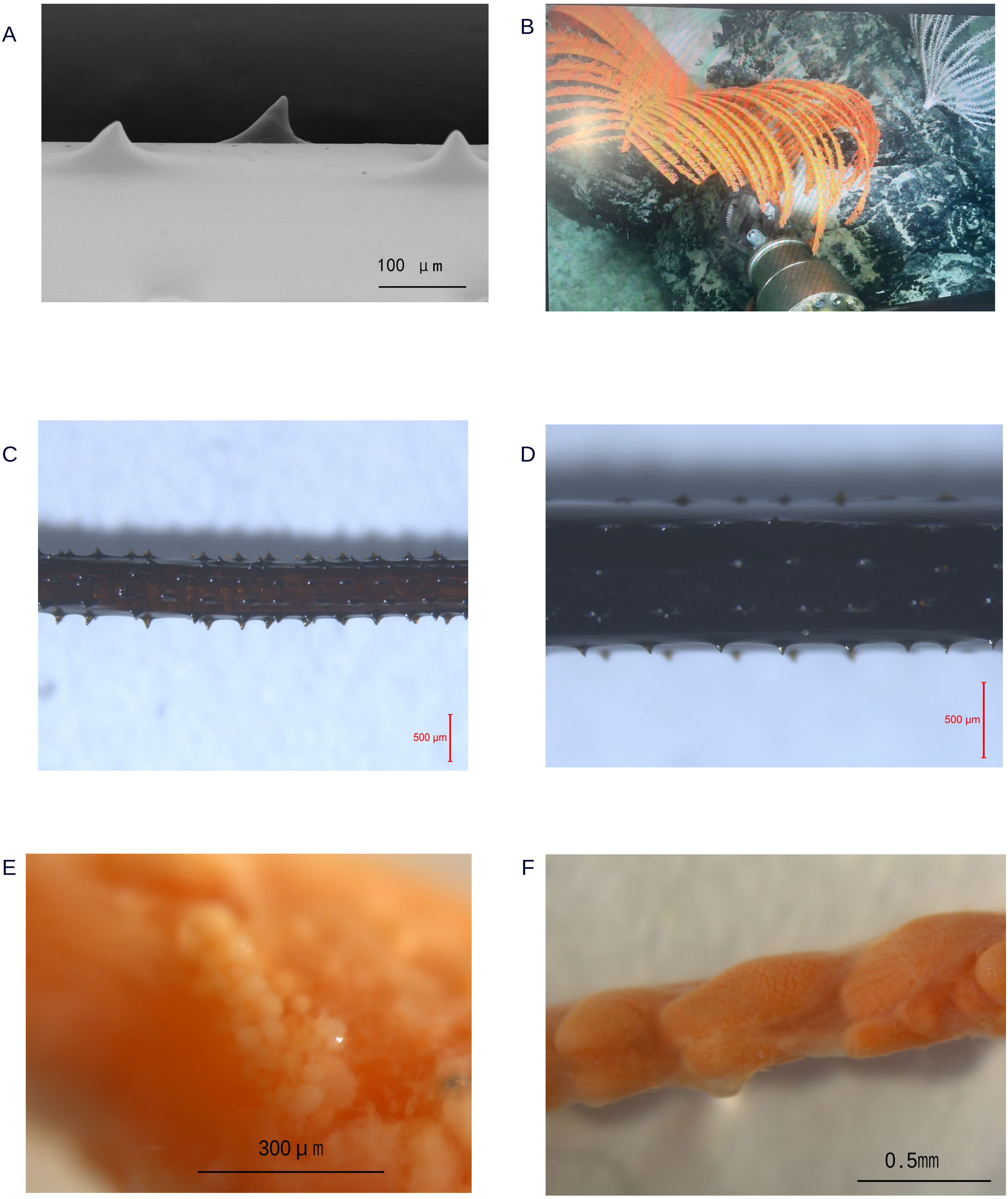


**Fig. S3.** Morphology of C1253.(A) Electron micrograph, (B) in situ photograph, (C-F) optical photograph.

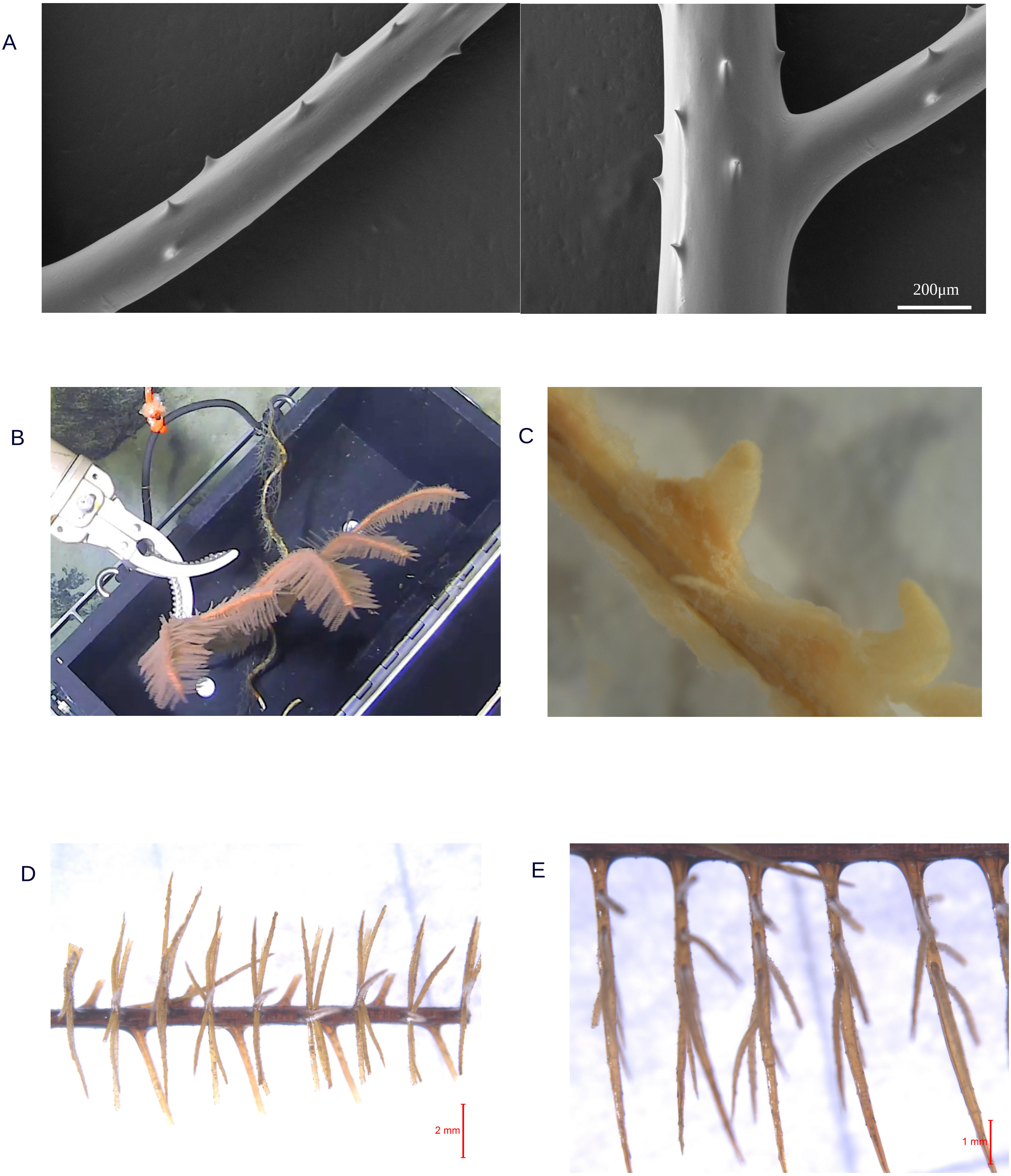


**Fig. S4.** Morphology of C1180 (A) Electron micrograph, (B) in situ photograph, (C-E) optical photograph

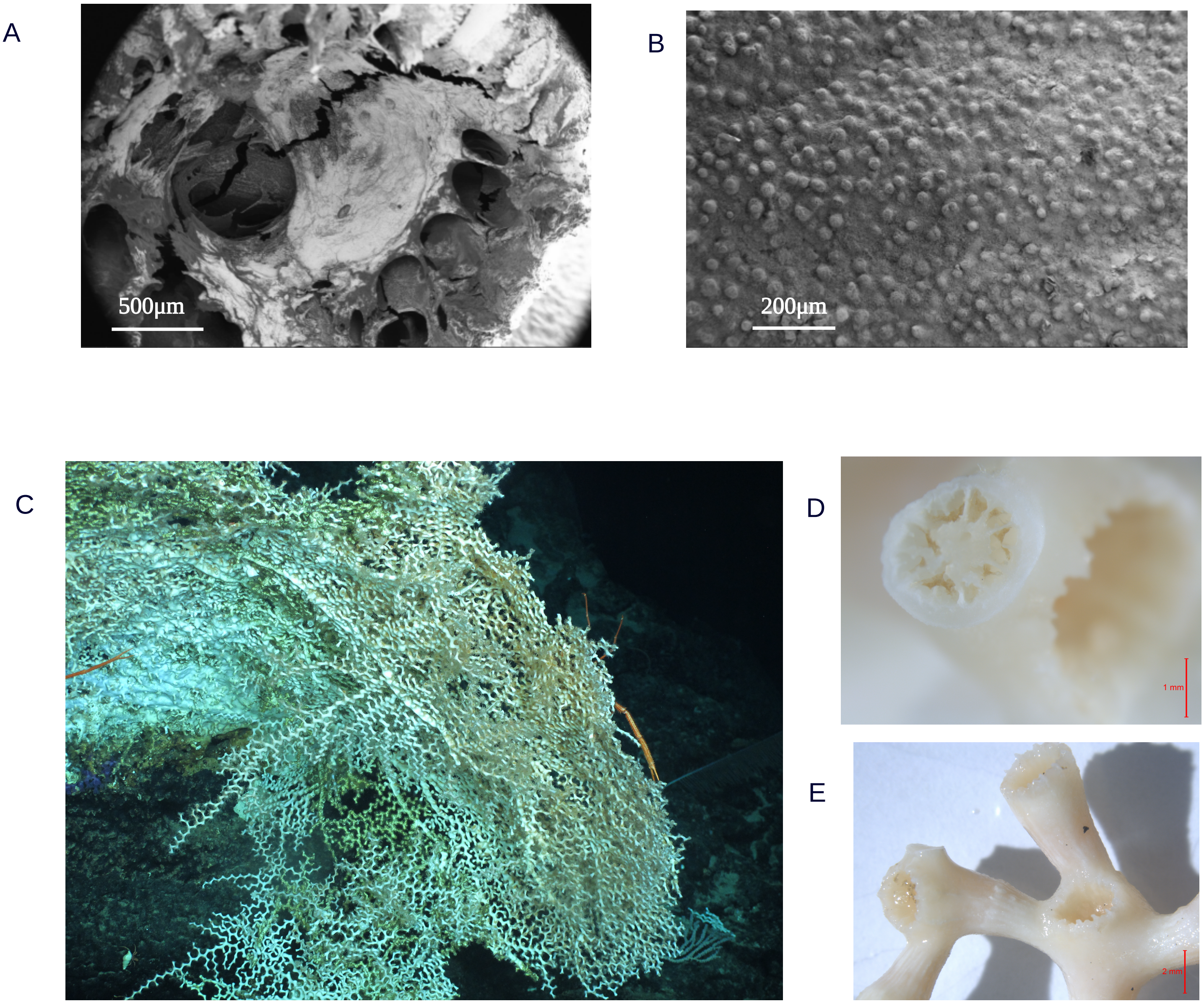


**Fig. S5.** Morphology of C1392 (A, B) Electron micrograph, (C) in situ photograph, (D, E) optical photograph

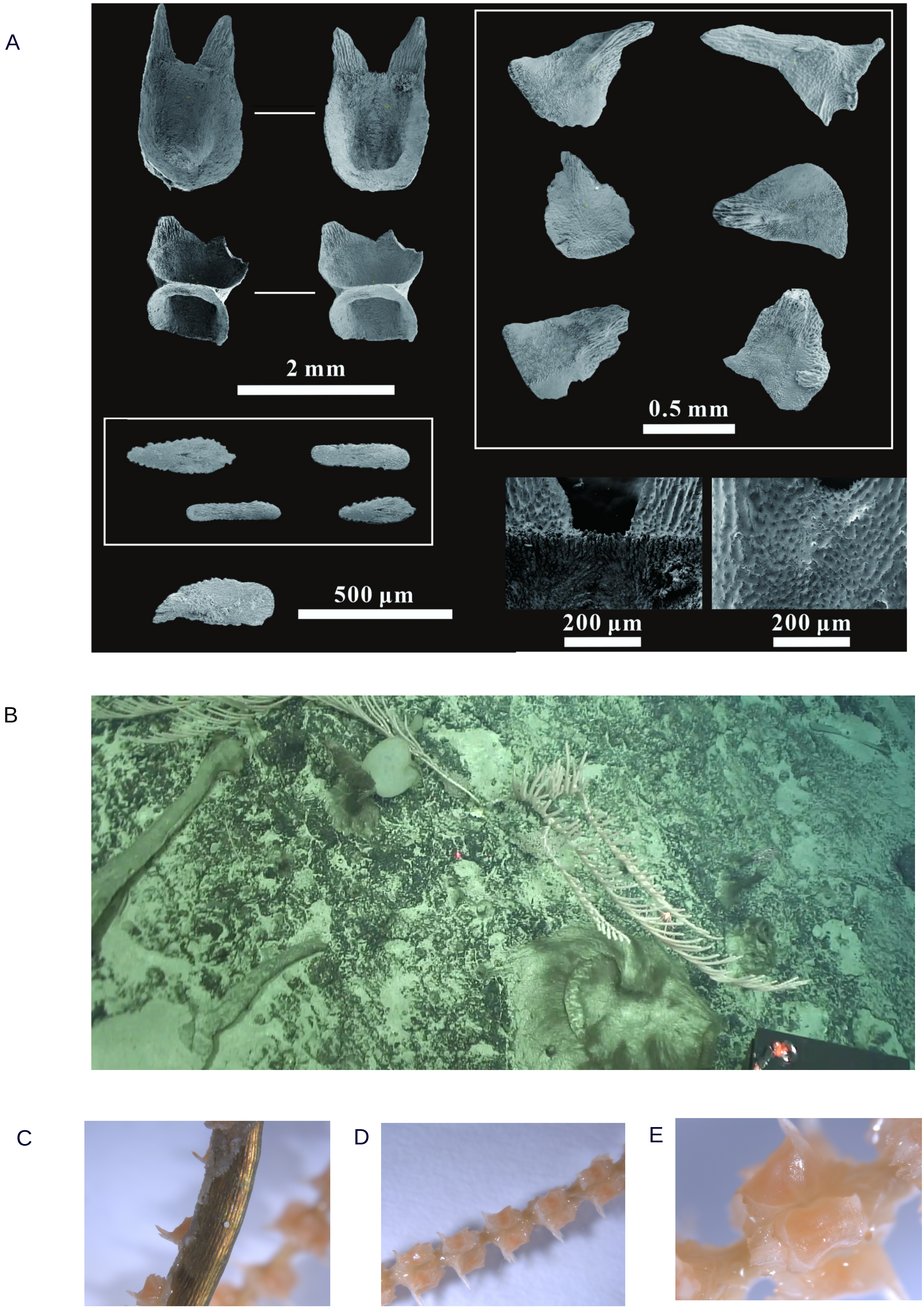


**Fig. S6.** Morphology of  C1633 .(A) Electron micrograph, (B) in situ photograph, (C-E) optical micrograph

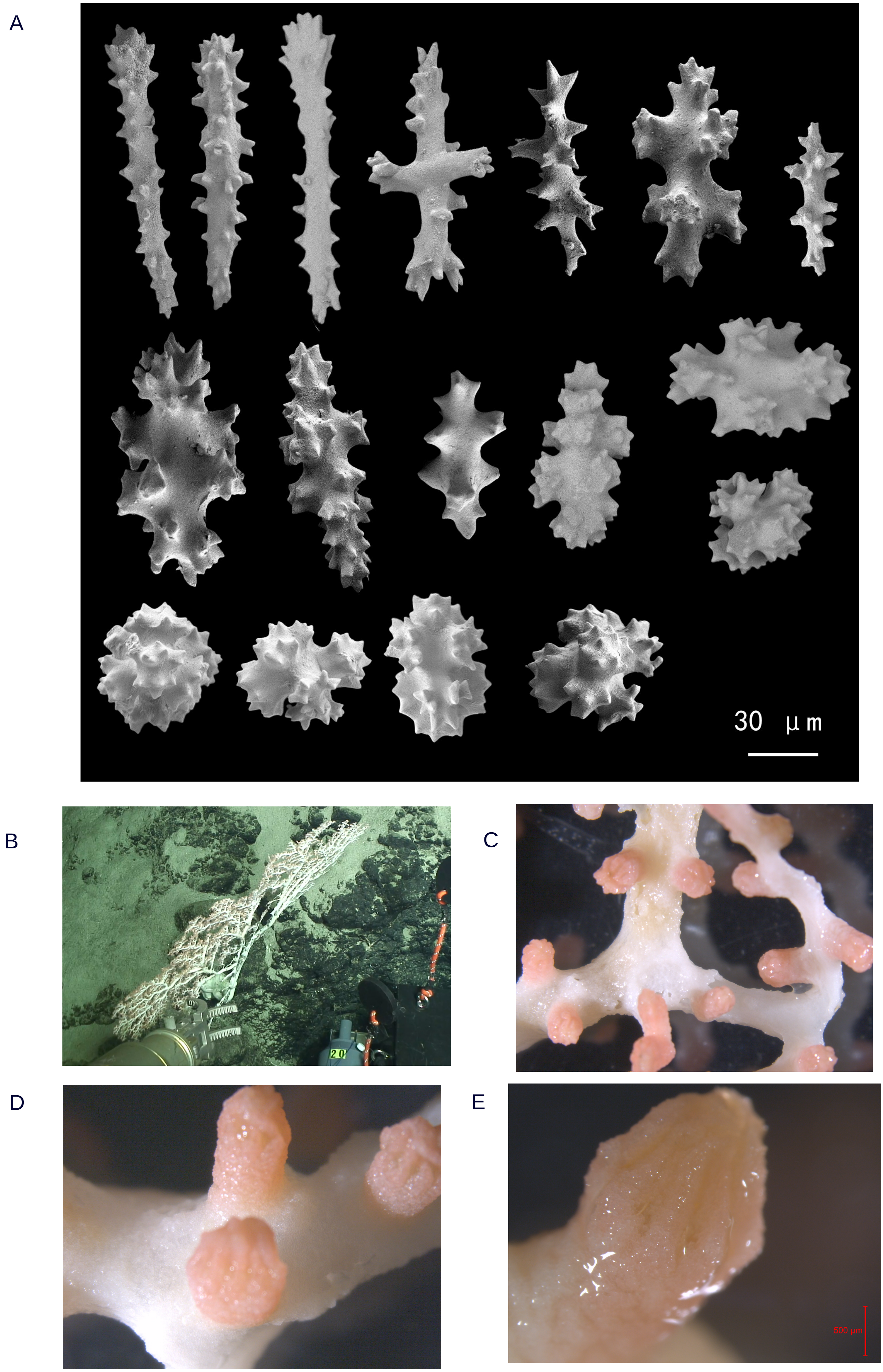


**Fig. S7.** Morphology of  C0925 (A) Electron micrograph, (B) in situ photograph, (C-E) optical photograph.

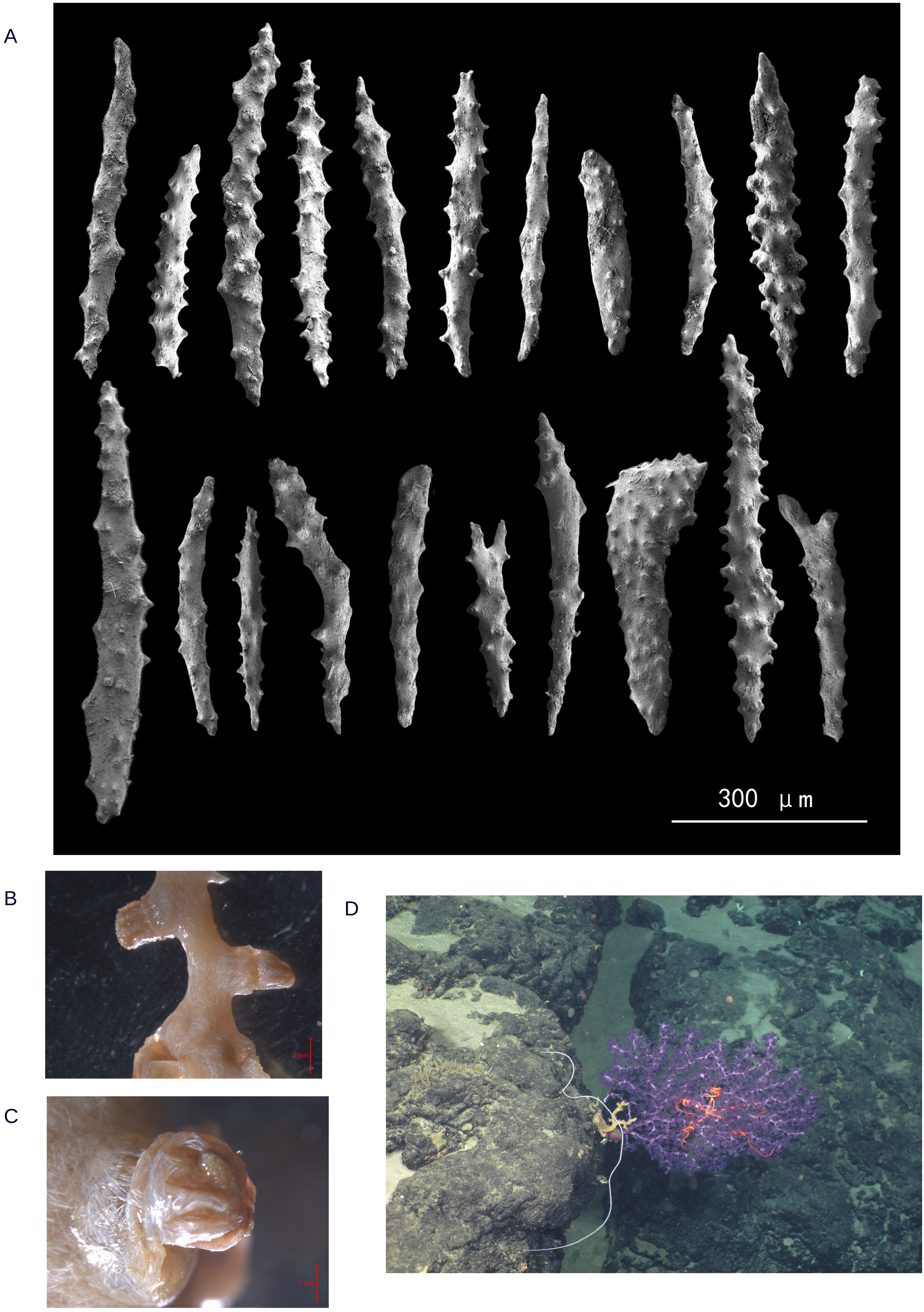


**Fig. S8.** Morphology of C1329 (A) Electron micrograph, (B, C) optical photograph, (D) in situ photograph.

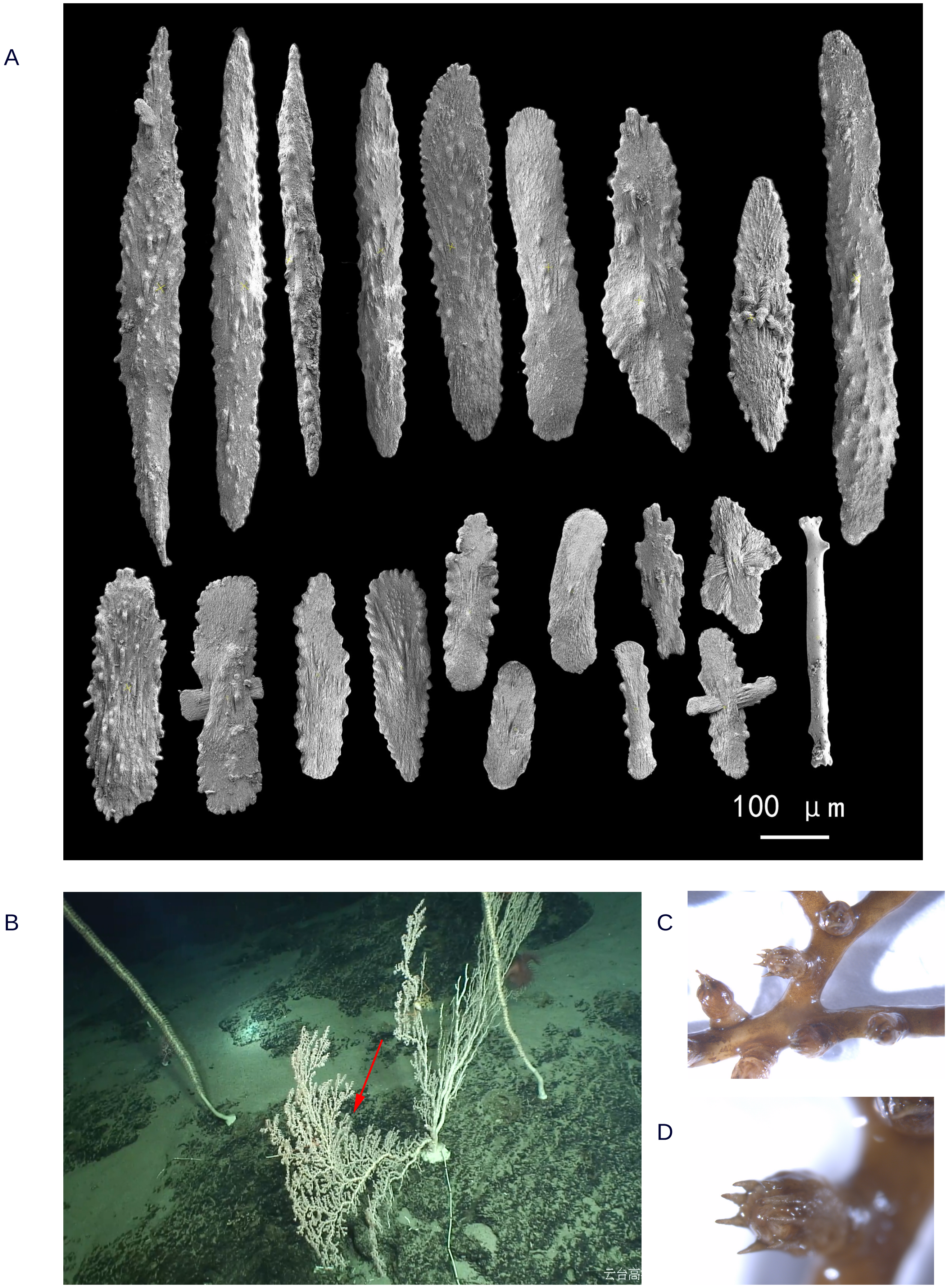


**Fig. S9.** Morphology of  C1356 (A) Electron micrograph, (B) in situ photograph, (C-D) optical photograph.

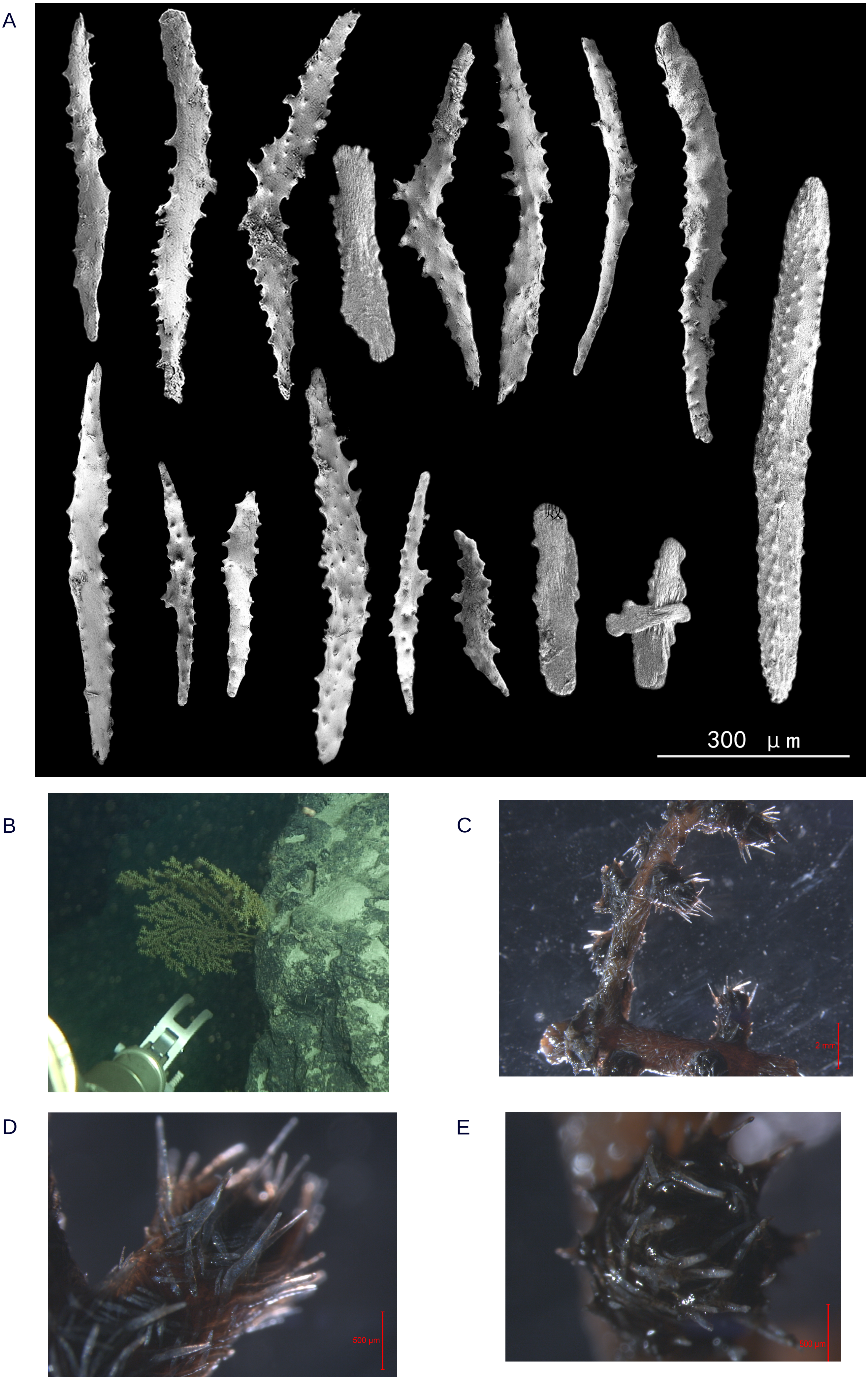


**Fig. S10.** Morphology of C1440. (A)Electron micrograph, (B) in situ photograph, (C, D, E) optical photograph.

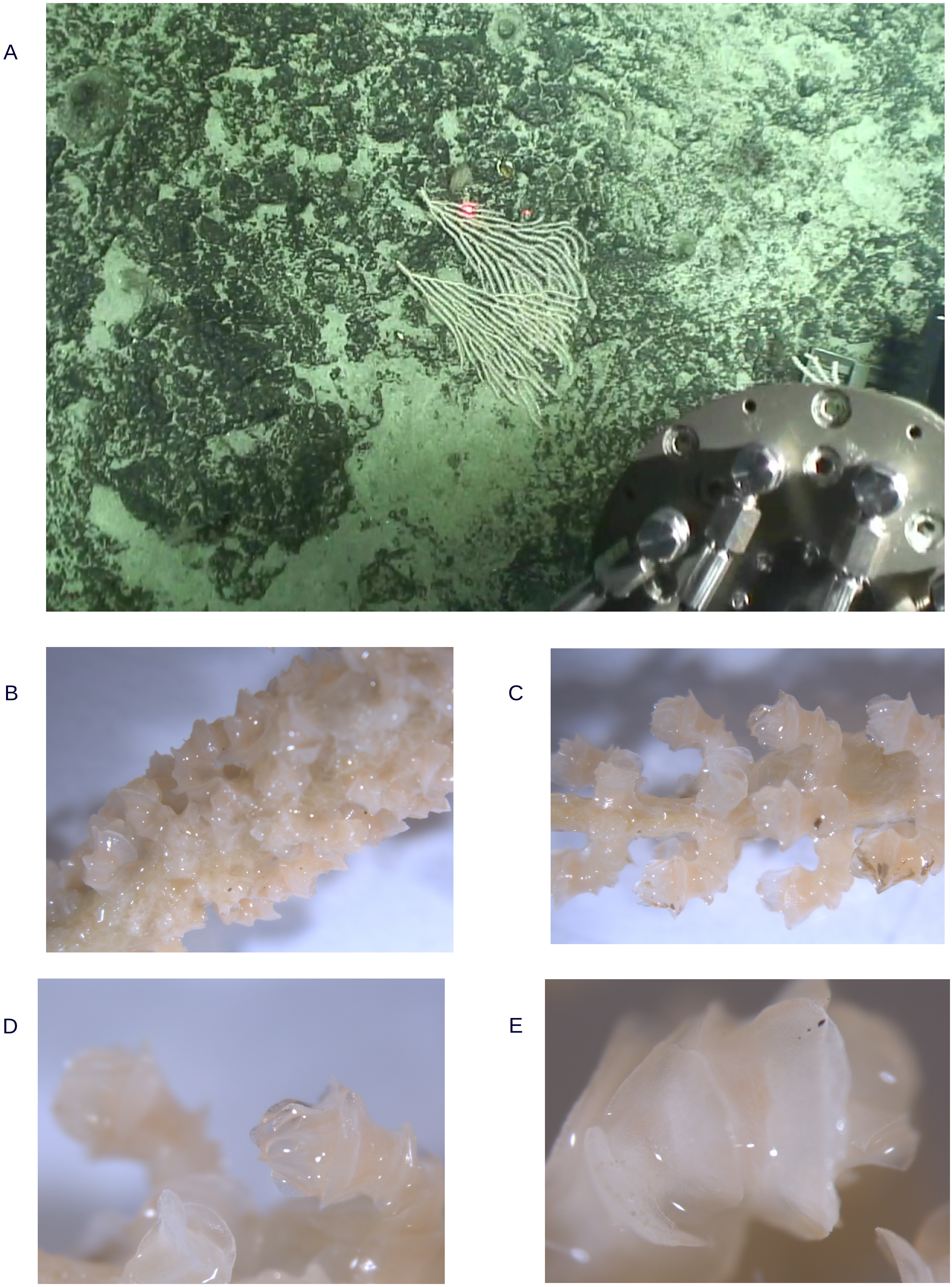


**Fig. S11.** Morphology of  C1773 (A) in situ photograph, (B-E) optical photograph

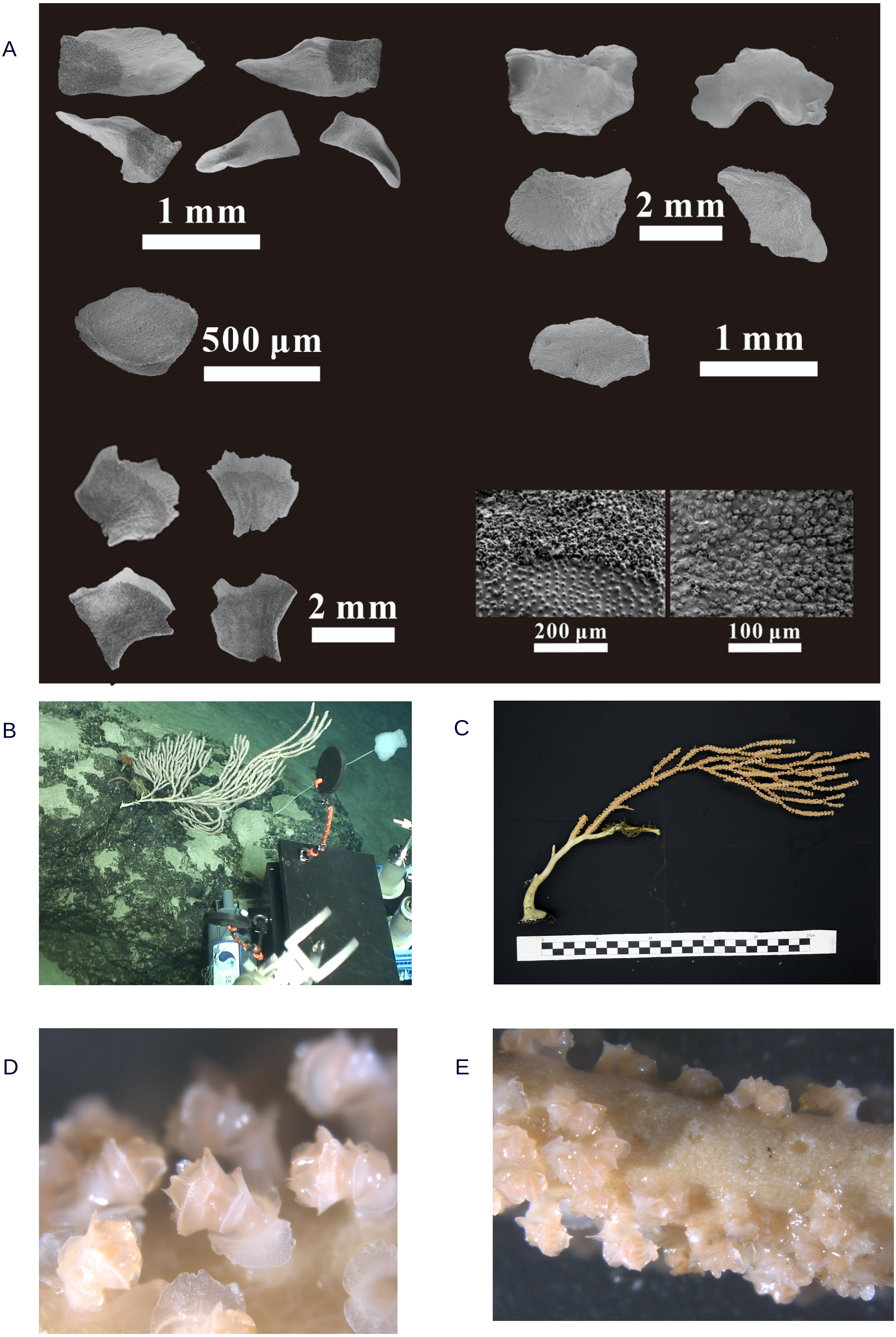


**Fig. S12.** Morphology of C0905 (A) Electron micrograph, (B) in situ photograph, (C-E) optical photograph.

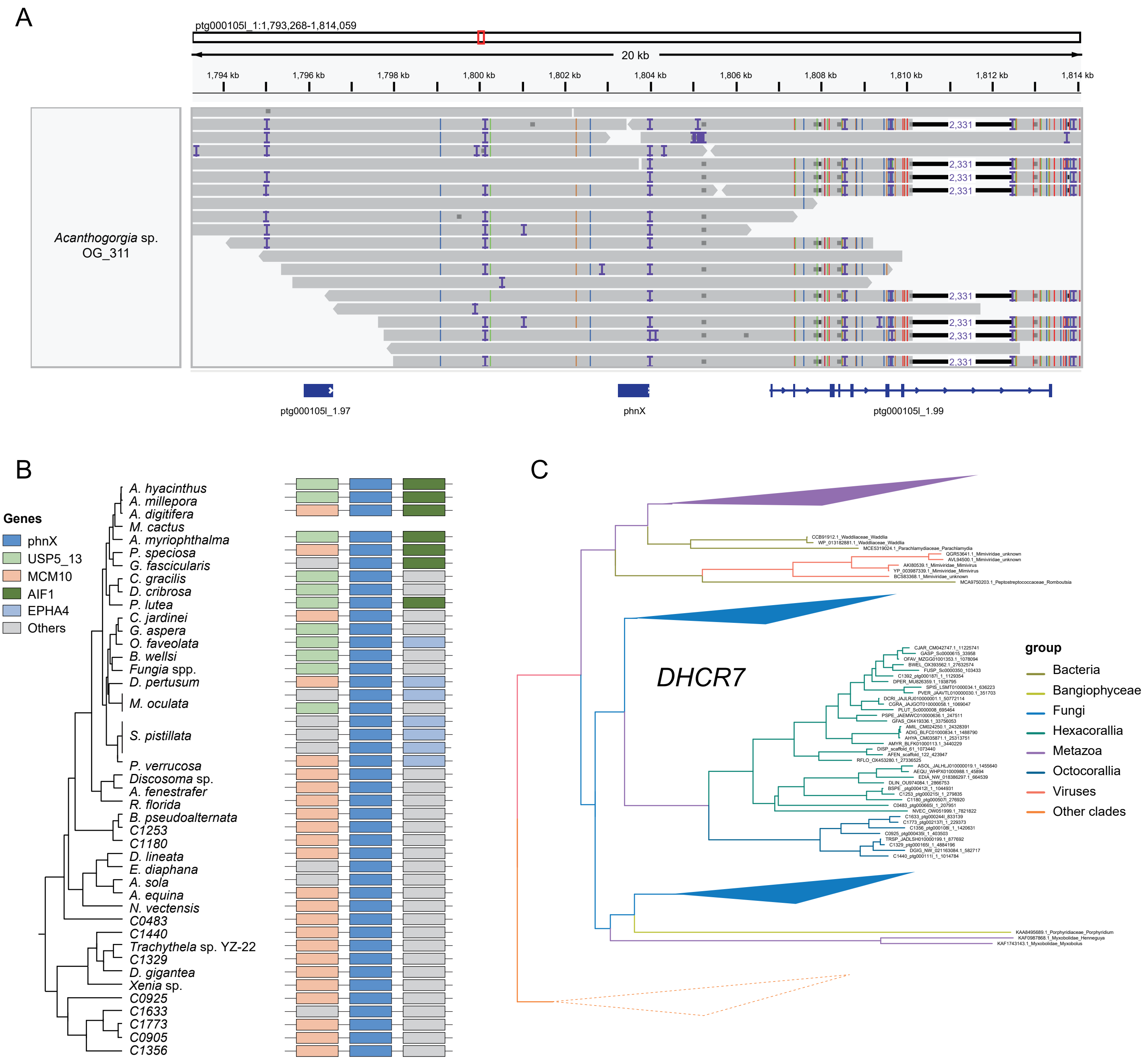


**Fig. S13.** Phylogenetic tree, upstream and downstream genes and verification of horizontal transferred genes. (A) Genome coverage of PacBio CCS reads near the *phnX* gene in C1440 (B) Genes upstream and downstream of the coral *phnX* gene. (C) Phylogenetic tree of the *DHCR7* gene. Branches containing the same taxonomy or are too far away from the coral lineage are collapsed.

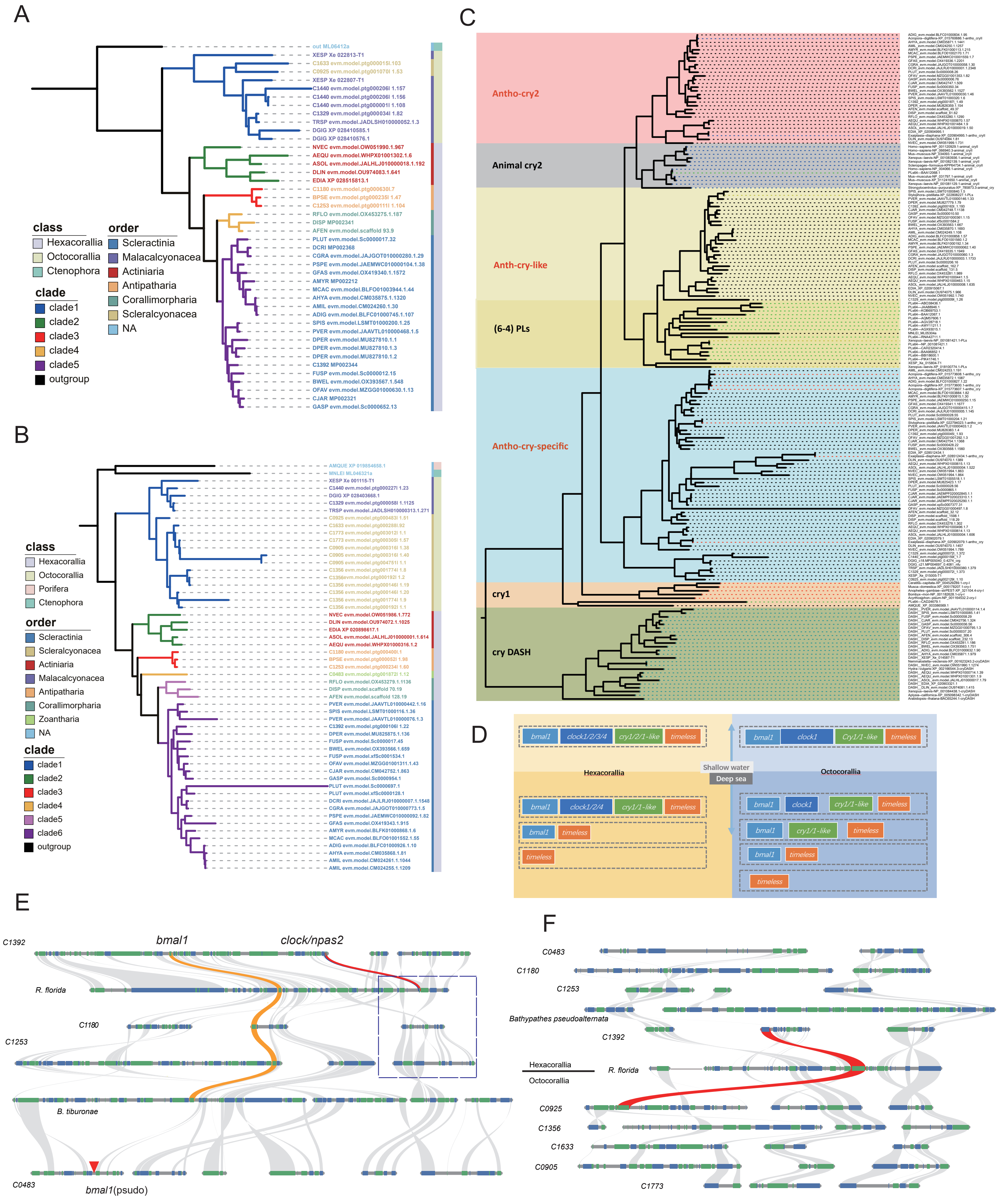
A
class
Hexacorallia
Octocorallia
Ctenophora
order
Scleractinia
Malacalcyonacea
Actiniaria
Antipatharia
Corallimorpharia
Scleralcyonacea
NA
clade
clade1
clade2
clade3
clade4
clade5
outgroup
B
class
Hexacorallia
Octocorallia
Porifera
Ctenophora
order
Scleractinia
Scleralcyonacea
Actiniaria
Malacalcyonacea
Antipatharia
Corallimorpharia
Zoantharia
NA
clade
clade1
clade2
clade3
clade4
clade5
clade6
outgroup
C
Antho-cry2
Animal cry2
Anth-cry-like
(6-4) PLs
Antho-cry-specific
cry1
cry DASH
D
bmal1
clock1/2/3/4
cry1/2/1-like
timeless
Hexacorallia
bmal1
clock1/2/4
cry1/1-like
timeless
bmal1
timeless
timeless
Shallow water
Deep sea
bmal1
clock1
Cry1/1-like
timeless
Octocorallia
bmal1
clock1
cry1/1-like
timeless
bmal1
cry1/1-like
timeless
bmal1
timeless
timeless
E
C1392
bmal1
clock/npas2
R. florida
C1180
C1253
B. tiburonae
C0483
bmal1(psudo)
F
C0483
C1180
C1253
Bathypathes pseudoalternata
C1392
Hexacorallia
Octocorallia
R. florida
C0925
C1356
C1633
C0905
C1773

**Fig. S14.** The phylogenetic and synteny of bmal1, cry, and timeless genes. (A, B, C) Phylogenetic tree of the genes bmal1, timeless, and cry. (D) Overview of key biological rhythm genes of shallow and deep-sea corals. (E, F) Synteny of bmal1 and clock genes among deep-sea corals. The red triangle of C0483 in (E) indicates pseudogenization. Blues boxes indicate lost regions in deep-sea corals.

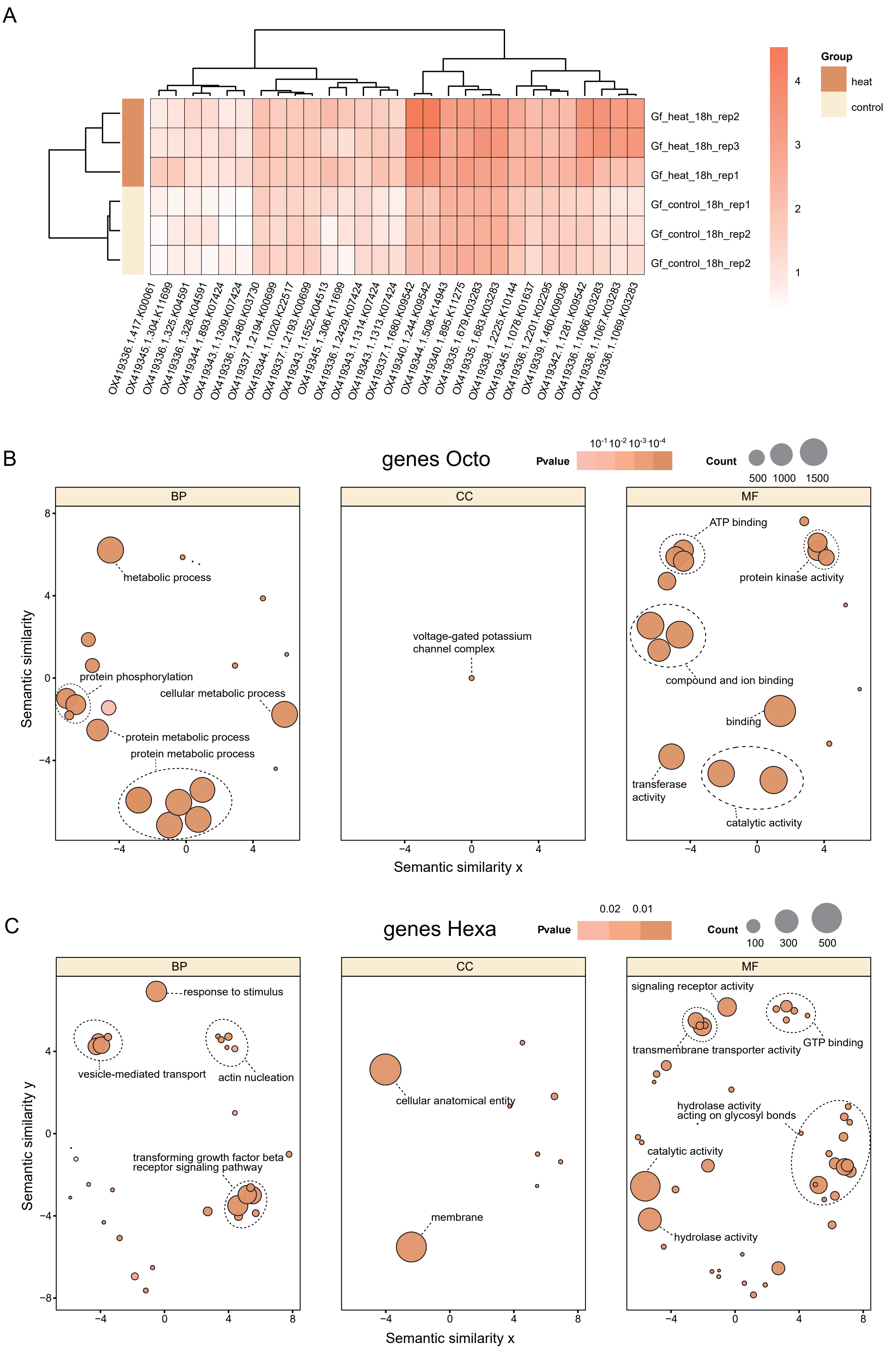
A
Group
heat
control
Gf_heat_18h_rep2
Gf_heat_18h_rep3
Gf_heat_18h_rep1
Gf_control_18h_rep1
Gf_control_18h_rep2
Gf_control_18h_rep2
OX419336.1.417.K00061
OX419345.1.304.K11699
OX419336.1.325.K04591
OX419336.1.328.K04591
OX419344.1.893.K07424
OX419343.1.1309.K07424
OX419336.1.2480.K03730
OX419337.1.2194.K00699
OX419344.1.1020.K22517
OX419337.1.2193.K00699
OX419343.1.1552.K04513
OX419345.1.306.K11699
OX419336.1.2429.K07424
OX419343.1.1314.K07424
OX419343.1.1313.K07424
OX419337.1.1680.K09542
OX419340.1.244.K09542
OX419344.1.508.K14943
OX419340.1.895.K11275
OX419335.1.679.K03283
OX419335.1.683.K03283
OX419338.1.2225.K10144
OX419345.1.1078.K01637
OX419336.1.2201.K02295
OX419339.1.460.K09036
OX419342.1.1281.K09542
OX419336.1.1066.K03283
OX419336.1.1067.K03283
OX419336.1.1069.K03283
B
genes Octo
Pvalue
Count
500 1000 1500
BP
CC
MF
Semantic similarity
Semantic similarity x
metabolic process
protein phosphorylation
cellular metabolic process
protein metabolic process
protein metabolic process
voltage-gated potassium channel complex
ATP binding
protein kinase activity
compound and ion binding
binding
transferase activity
catalytic activity
C
genes Hexa
Pvalue
0.02 0.01
Count
100 300 500
Semantic similarity y
Semantic similarity x
response to stimulus
vesicle-mediated transport
actin nucleation
transforming growth factor beta receptor signaling pathway
cellular anatomical entity
membrane
signaling receptor activity
transmembrane transporter activity
GTP binding
hydrolase activity acting on glycosyl bonds
catalytic activity
hydrolase activity

**Fig. S15.** Expression heat map of genes differentially expressed in response to heat by the stone coral Galaxea fascicularis and function of different gene families in Hexacorallia and Octocorallia deep-sea corals. (A) Normalized gene expression of genes upregulated under heat conditions by G. fascicularis. (B) Enriched GO function of genes that expanded in Octocorallia deep-sea corals. (C) Enriched GO function of genes that expanded in Hexacorallia deep-sea corals.
